# Harnessing natural modularity of cellular metabolism to design a modular chassis cell for a diverse class of products by using goal attainment optimization

**DOI:** 10.1101/748350

**Authors:** Sergio Garcia, Cong T. Trinh

**Affiliations:** Department of Chemical and Biomolecular Engineering, The University of Tennessee, Knoxville, TN, United States; Center for Bioenergy Innovation, Oak Ridge National Laboratory Oak Ridge, TN, United States

**Keywords:** Biocatalysis, modular cell, ModCell, modular design, metabolic network modeling, constraint-based modeling, multi-objective optimization, mixed integer linear programming, goal programming, Benders decomposition

## Abstract

Living cells optimize their fitness against constantly changing environments to survive. Goal attainment optimization is a mathematical framework to describe the simultaneous optimization of multiple conflicting objectives that must all reach a performance above a threshold or goal. In this study, we applied goal attainment optimization to harness natural modularity of cellular metabolism to design a modular chassis cell for optimal production of a diverse class of products, where each goal corresponds to the minimum biosynthesis requirements (e.g., yields and rates) of a target product. This modular cell design approach enables rapid generation of optimal production strains that can be assembled from a modular cell and various exchangeable production modules and hence accelerates the prohibitively slow and costly strain design process. We formulated the modular cell design problem as a blended or goal attainment mixed integer linear program, using mass-balance metabolic models as biological constraints. By applying the modular cell design framework for a genome-scale metabolic model of *Escherichia coli*, we demonstrated that a library of biochemically diverse products could be effectively synthesized at high yields and rates from a modular (chassis) cell with only a few genetic manipulations. Flux analysis revealed this broad modularity phenotype is supported by the natural modularity and flexible flux capacity of core metabolic pathways. Overall, we envision the developed modular cell design framework provides a powerful tool for synthetic biology and metabolic engineering applications such as industrial biocatalysis to effectively produce fuels, chemicals, and therapeutics from renewable and sustainable feedstocks, bioremediation, and biosensing.

## 1 Introduction

Microbial metabolism can be engineered to produce a large space of molecules from renewable and sustainable feedstocks.^1^ Currently, only a handful of fuels and chemicals out of the many possible molecules offered by nature are industrially produced by microbial conversion, mainly because the strain engineering process is too laborious and expensive.^2^ Thus, innovative technologies enabling rapid and economically-feasible strain engineering are needed to harness a large space of industrially-relevant biochemicals.^1–3^ To tackle this challenge, the principles of modular design that have shown great success in traditional engineering disciplines can be adapted to construct modular cell biocatalysts in a plug-and-play fashion with minimal strain optimization cycles.^4^

Multi-objective optimization is a powerful mathematical framework widely applied in engineering disciplines to tackle the optimal design of a complex system with multiple conflicting objectives.^5,6^ This framework has recently been exploited for not only explaining the modularity of natural biological systems that enable cellular robustness and adaptability^7–11^ but also implementing modular engineering design.^12^ Using multi-objective optimization, microbial metabolism can be redirected to generate modular production strains that are systematically assembled from an engineered modular cell and exchangeable production modules, each of which synthesizes a target molecule.^13^ This modular cell (ModCell) design approach, known as ModCell2, uses the principles of mass balance and thermodynamics of biochemical reaction networks to predict metabolic fluxes upon genetic manipulations.^13,14^ Based on such flux predictions, a multi-objective optimization problem is then formulated and solved with a multi-objective evolutionary algorithm (MOEA)^15,16^ to yield a sample of the Pareto front (i.e., the set of optimal solutions to the problem with minimal trade-offs among objectives) that a designer can explore genetic manipulation targets for modular cell engineering.

In this study, we developed ModCell2-MILP, a ModCell2-based formulation to be compatible with mixed integer linear programming (MILP) algorithms. This framework presents a significant advancement from ModCell2 in solving the multi-objective strain design problem for modular cell engineering. Specifically, ModCell2-MILP is developed to (i) guarantee optimal solutions, (ii) completely enumerate alternative solutions of a target design, and (iii) describe practical engineering goals more directly (e.g., design of a modular cell where all production modules lead to a product yield above 50% of the theoretical maximum). By applying ModCell2-MILP to analyze the genome-scale metabolic network of *Escherichia coli*, we could identify a universal modular cell that is compatible with a diverse class of production modules. Finally, we shed light on the underlying features of the universal modularity phenotype by systematically analyzing feasible flux distributions of all modular production strains. We anticipate ModCell2-MILP can provide a powerful tool for not only elucidating natural and synthetic metabolic modularity but also rationally designing modular production strains for novel synthetic biology and metabolic engineering applications.

## 2 Methods

### 2.1 Modular cell design

#### 2.1.1 Design principles

ModCell design enables rapid assembly of production strains with desirable phenotypes from a modular (chassis) cell.^4,13,17^ More specifically, a modular cell contains core metabolic pathways shared among production modules (Figure 1a). The chassis interfaces with the modules through enzymatic and genetic synthesis machinery and precursor metabolites (Figure 1b). Modules contain auxiliary regulatory and metabolic pathways (Figure 1c) that enable a desired phenotype for optimal biosynthesis of a target molecule, for example, weak growth coupled to product formation (*wGCP*), where a positive correlation between growth and product synthesis rates is enforced (Figure 1d).^13,18,19^ The design objective phenotypes are determined from cellular growth and product synthesis rates based on steady-state, mass-balance metabolic models.^20^ A modular cell is said to be *compatible* with a module if the design objective of the resulting production strain is above a specified threshold. The different biochemical nature of production modules to synthesize target metabolites can make the design objectives compete with each other and also the cellular objectives (e.g., biomass formation) compete with the engineering objectives (e.g., product formation), turning the ModCell design problem into a multi-objective and multi-level optimization problem.

**Figure 1:**
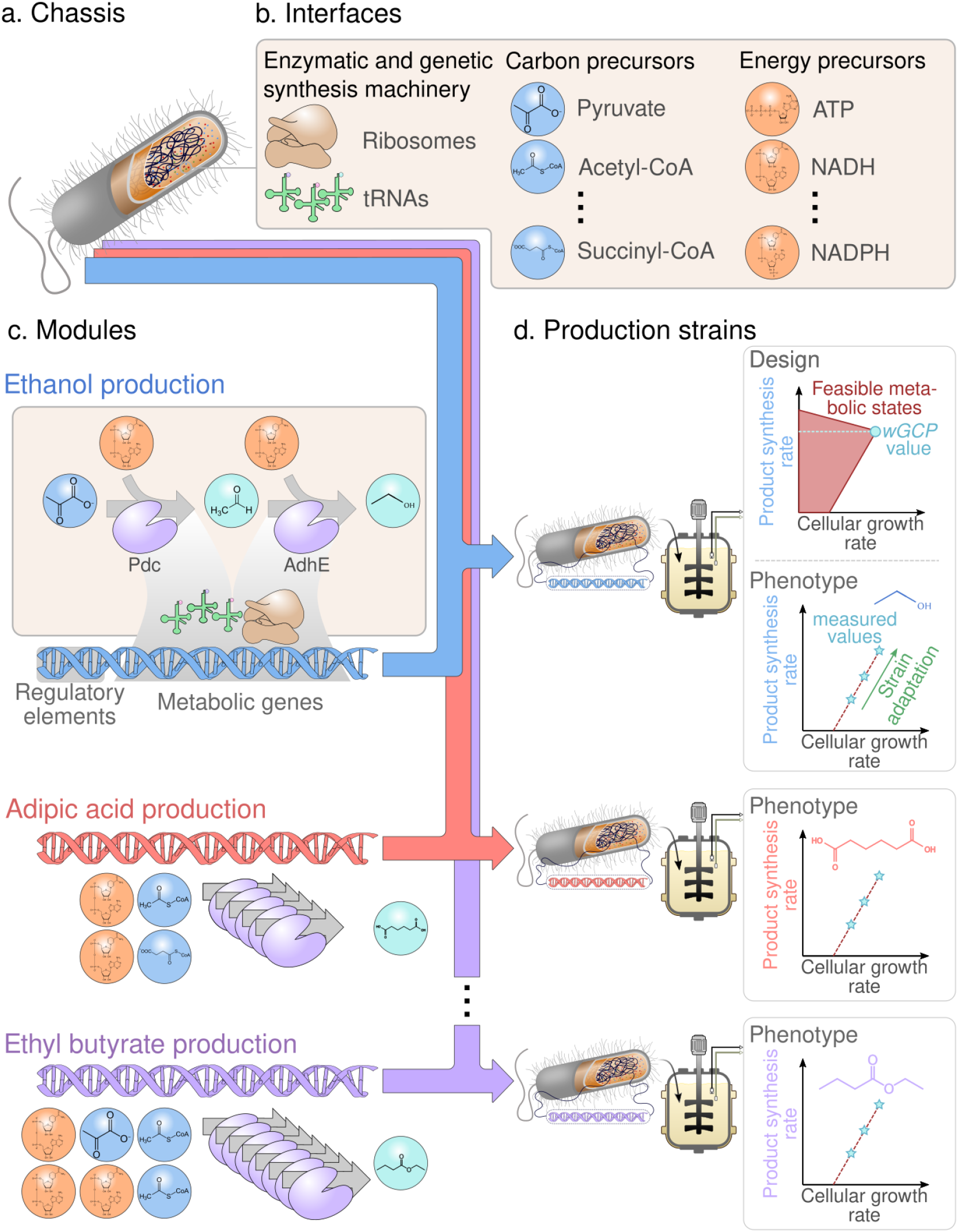
Principles of modular cell design. (a) Modular cell chassis. (b) Interfaces. (c) Production modules. (d) Production strains. A modular cell is designed to provide the necessary precursors for biosynthesis pathway modules that are independently assembled with the modular cell to generate production strains exhibiting desirable phenotypes. The *wGCP* phenotype, one of the possible design objectives, enforces the coupling between the desirable product synthesis rate and the maximum cellular growth rate.

#### 2.1.2 Multi-objective optimization formulation

The modular cell design problem is stated as a general multi-objective optimization problem of the form:

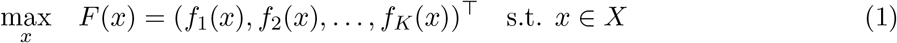

where *f_k_* is the desirable phenotype for production module *k*, *x* are the problem variables including binary design variables corresponding to genetic manipulations, and *X* is the set of constraints including mass balance of metabolism. Optimal solutions for the multi-objective optimization problem (1) are defined using the concept of domination: A vector *a* = (*a*_1_,…, *a_K_*)^⊤^ *dominates* another vector *b* = (*b*_1_,…, *b_K_*)^⊤^, denoted as *a* ≺ *b*, if and only if *a_i_* ≥ *b_i_* ∀*i* ∈ {1,2,…, *K*} and *a_i_* ≠ *b_i_* for at least one *i*. A feasible solution *x*\* ∈ *X* of the multi-objective optimization problem is called a Pareto optimal solution if and only if there does not exist a vector *x′* ∈ *X* such that *F*(*x*\*) ≺ *F*(*x*\*). The set of all Pareto optimal solutions is called Pareto set:

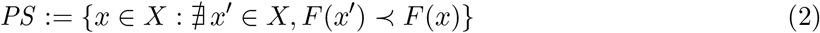

The projection of the Pareto set in the objective space is denoted as Pareto front:

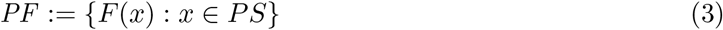

Different feasible points in *PS* (i.e., different genetic manipulations) which map to a single point in *PF* (i.e., the same phenotype) are denoted *alternative solutions*.

The design variables *x* in ModCell2 correspond to chassis reaction deletions, that remove undesired metabolic functions, and module reaction insertions, that allow to identify optimal module configurations without extensive prior knowledge of the product synthesis pathway. The constraint set *X* is comprised of two types: (i) flux simulation constraints (e.g., mass balance, reaction reversibility, and flux bound) that allow to predict fluxes in the design objectives upon genetic manipulations, and (ii) implementation constraints that involve the maximum number of reaction deletions in the chassis (denoted by *α*) and the maximum number of module reaction insertions per module (denoted by *β*). The following sections describe the problem formulation in detail using the definitions compiled in Section 5.

#### 2.1.3 Design objectives

Design objectives, *f*_k_, that correspond to specific metabolic phenotypes within the space of feasible steady-state reaction fluxes, Π*_km_*, of production network *k* (i.e., the combination of the chassis network with the production module *k*) and metabolic state *m*, are defined as follows:

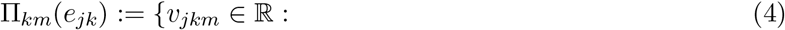

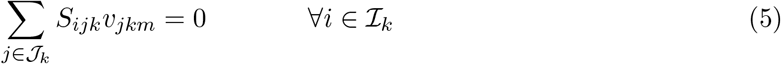

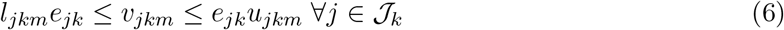

Here, *v_jkm_* is the rate (mmol/gCDW/hr) of reaction *j* in production network *k* under metabolic state *m*. Constraint (5) enforces mass balance for all metabolites according to reaction stoichiometry given by the coefficients *S_ijk_*, and constraint (6) imposes bounds, *l_jkm_* and *u_jkm_*, for the metabolic fluxes according to reaction reversibility, experimentally measured values, and specified metabolic state. The binary variable *e_jk_* is used in the overall optimization problem to indicate whether reaction *j* in production network *k* is removed and thus cannot carry any flux. Two metabolic states m are considered, growth and non-growth, denoted *μ* and 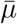, respectively. These states are differentiated by their flux bounds *l_jkm_* and *u_jkm_*. For growth state, the lower bound of the biomass formation reaction that represents cell division, *v_Xkm_*, is set to a minimum value of *γ*, i.e., 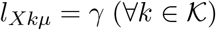, while there is no upper limit to growth, i.e., 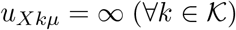. On the other hand, for the non-growth state both bounds are set to 0, i.e., 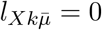 and 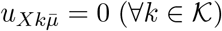.

Given the feasible metabolic flux space, Π*_km_*, the following design objectives, based on the product synthesis rate reaction, *v_Pkm_*, are of interest:

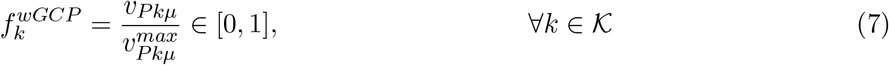

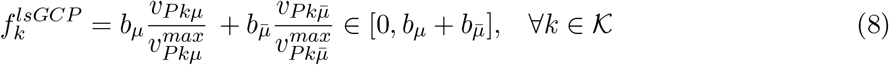

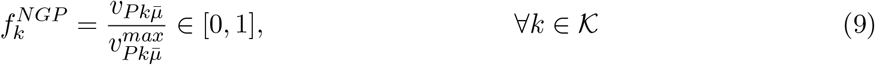

The product synthesis fluxes, including 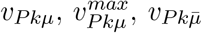, and 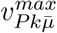, are computed by solving the following linear programming problems:

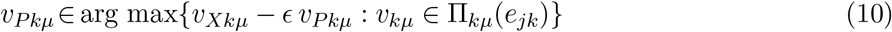

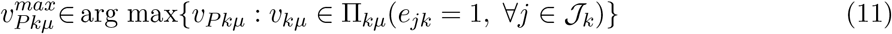

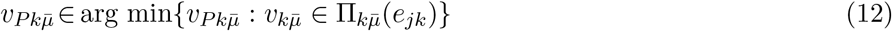

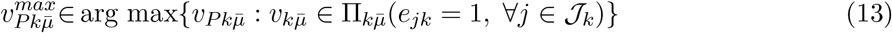

The maximum product synthesis fluxes (11) and (13) used for objective scaling are only calculated once by not using any deleted reactions (*e_jk_* = 1), while the target phenotype fluxes (10) and (12) are functions of the deleted reactions *e_jk_*. The design objectives, *wGCP* (7), *lsGCP* (8), and *NGP* (9), were previously proposed^13^ and briefly described here. The weak growth coupled to product formation objective (*wGCP*) (7) seeks to maximize the minimum product rate at the maximum cellular growth, which is accomplished by a titled objective function^21^ (10). The linearized strong growth coupled to product formation (*lsGCP*) (8) objective seeks to maximize the minimum product synthesis rate at the non-growth state 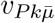 in addition to the goal of *wGCP*. Finally, the non-growth production (*NGP*) (9) objective seeks to optimize the minimum product synthesis rate during the non-growth state.

#### 2.1.4 Design constraints

All the constraints of the modular cell design problem are gathered as follows:

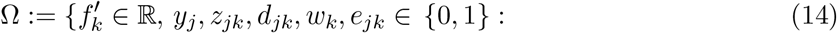

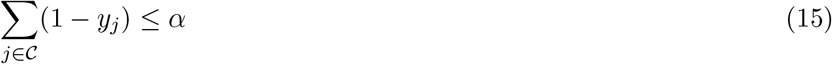

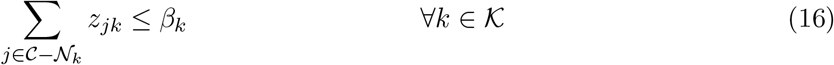

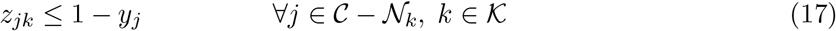

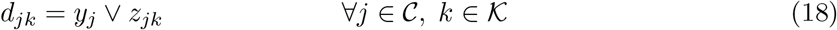

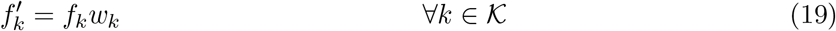

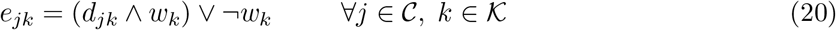

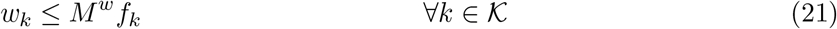

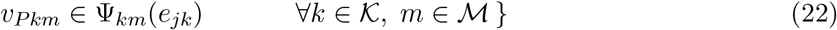

Constraints (15)-(18) are formulated for practical limitations and features of the modular cell. Specifically, the two variables that represent design choices for genetic manipulations include: (i) *y_j_* that takes a value of 0 if reaction *j* is deleted in the chassis (and consequently in all production networks) and 1 otherwise and (ii) *z_jk_* that takes a value of 1 if reaction *j* is inserted in production network *k*. The maximum number of reaction deletions, is limited by *α* through constraint (15) while the maximum number of module reactions in each module *β_k_* is imposed by (16). Constraint excludes non-candidate reactions 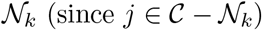 so that endogenous module reactions can be fixed (i.e., *z_jk_* = 1), according to problem-specific knowledge. Constraint (17) ensures that only reactions deleted in the chassis can be inserted back to the modules. Constraint (18) indicates that reaction *j* is deleted in production network *k* if the reaction is deleted in the chassis and not added as an endogenous module reaction. The designer can gradually increase *α* and *β_k_* to obtain solutions with higher performance.

Constraints (19)-(21) are introduced for modeling purposes. The indicator variable, *w_k_*, is introduced to allow for certain production networks to be ignored from the final solution. Without *w_k_*, the whole multi-objective problem becomes infeasible if a set of deletions renders one of the production networks infeasible (e.g., its minimum growth rate cannot be accomplished). However, in practice it is acceptable for some modules not to work with the chassis cell. If *w_k_* = 0, the objective value 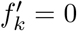 (19) and reaction deletions do not apply to network *k* since *e_jk_* = 1 (20); if *w_k_* = 1, 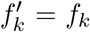 and *e_jk_* = *d_jk_*, where *f_k_* is any of the design objectives presented earlier (7)-(9). The use of *w_k_* is likely to introduce symmetry (i.e., alternative integer solutions with no practical meaning) due to cases where *f_k_* = 0 for a given *k* while the associated production network remains feasible, allowing *w_k_* to take a value of 0 or 1. This symmetry is removed by enforcing w¾ to be 0 if *f_k_* = 0 (21).

Finally, constraint (22) indicates that the fluxes featured in the design objectives, *v_Pkm_*, are contained in the polytope Φ*_km_*. The space of *v_Pkm_* is originally defined as an optimization problem (10)-(13), thus representing a non-linear constraint and turning the ModCell design problem into a bilevel optimization problem. These inner optimization problems are linearized, leading to Φ*_km_* as described in Section 2.1.6.

#### 2.1.5 Linearization of logical expressions

The logical expressions in Ω are replaced by the following linear constraints in the final problem formulation:

*d_jk_* = *y_j_* ∨ *z_jk_* corresponds to:

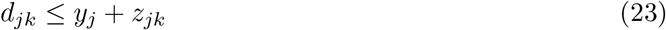

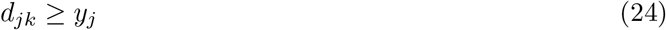

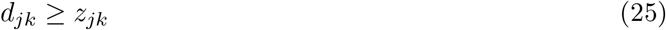

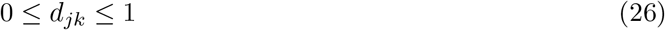
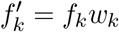 corresponds to:

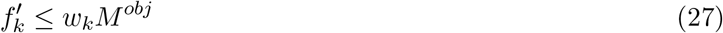

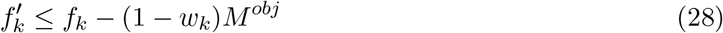

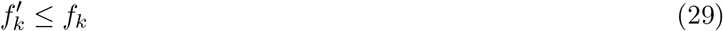

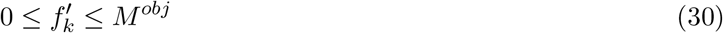
*e_jk_* = (*d_jk_* ∧ *w_k_*) ∨ ¬*w_k_*, given *r_jk_* = *d_jk_* ∧ *w_k_*, corresponds to:

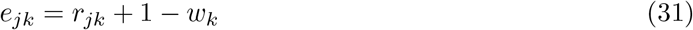

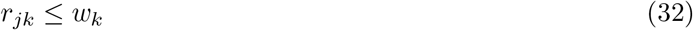

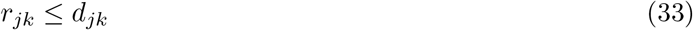

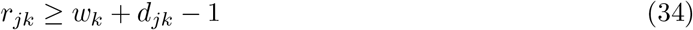

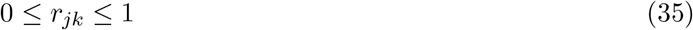

#### 2.1.6 Linearization of inner optimization problems

Non-linear constraints expressed as linear programming problems can be linearized using basic mathematical programming theory. Consider the following canonical linear program, with primal variables 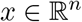 and its dual variables 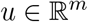:

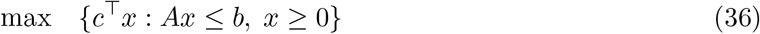

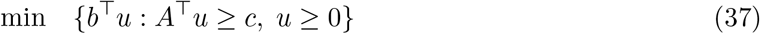

the strong duality theorem states that the objective functions of primal (36) and dual (37) are equal at their optima, *c*^⊤^ *x*\* = *b*^⊤^ *y*\*. Thus the optimal solution to the primal problem is described by the following linear constraints:

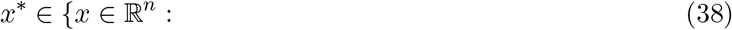

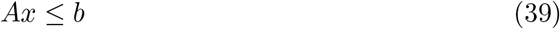

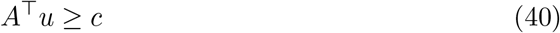

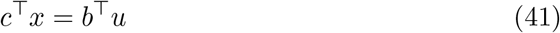

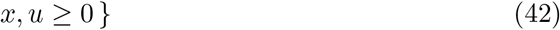

Using the strong duality theorem as presented by Maranas and Zomorrodi,^22^ the inner optimization problems (22) are linearized as follows:

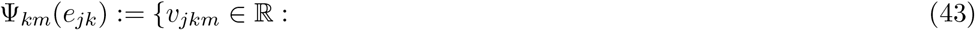

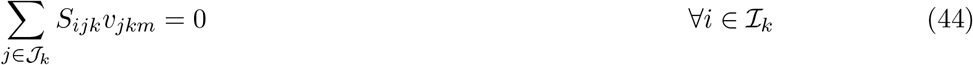

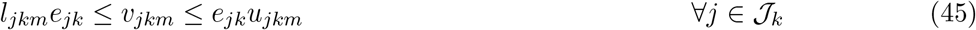

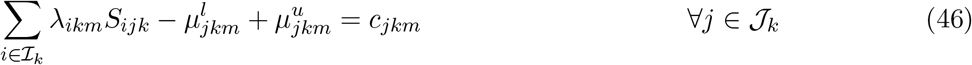

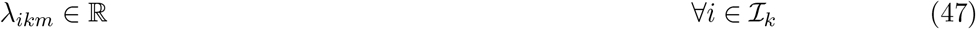

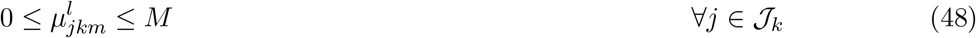

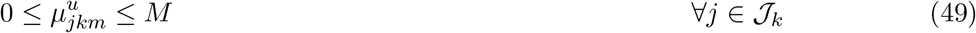

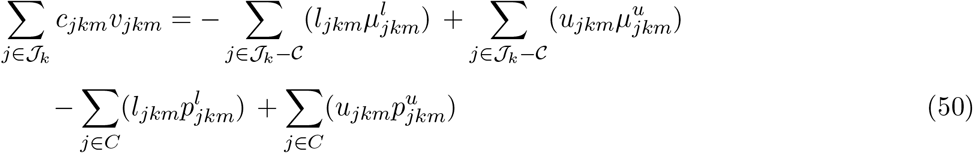

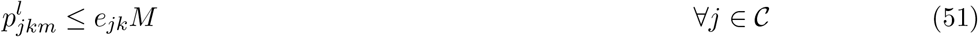

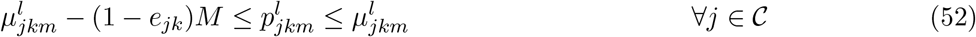

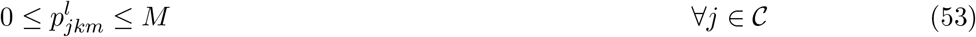

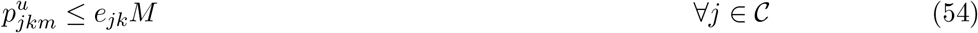

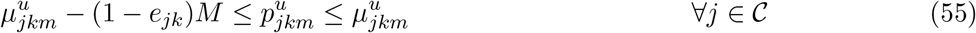

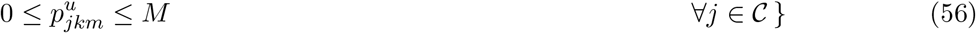

Constraints (44)-(45) correspond to the primal metabolic network problem and were introduced earlier in Π*_km_*. Constraints (46)-(49) correspond to the dual problem. We use the dual variables, λ*_ikm_*, for the primal mass balance constraints (44), together with 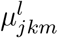 and 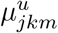 for the primal flux bound inequalities (45) involving lower and upper reaction bounds respectively. Constraints (47)-(49) emphasize the domain of the dual variables, with M being a large value above the expected value of any dual variable. Constraints (50)-(56) correspond to the strong duality equality. The left hand side of the strong duality equality (50) features the objectives presented in (10) for *m* = *μ* and (12) for 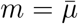. On the right hand side, products of binary and continuous variables appear, thus requiring linearization variables 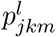 and 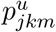. Constraints (51)-(56) ensure that 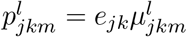 and 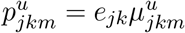.

#### 2.1.7 Conversion of a multi-objective problem into a single-objective problem

The multi-objective optimization problem (1) is now described entirely in terms of linear constraints through Ω. However, to make the formulation compatible with MILP solver algorithms, the objective function vector, *f*′, must be expressed as a scalar. To accomplish this without loss of relevant information, we employed blended and goal attainment formulations.

#### 2.1.8 Blended formulation

In the blended formulation,^23^ all objectives are summed as follows:

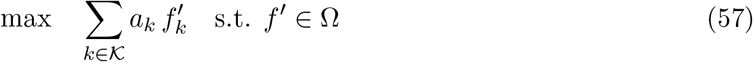

where *a_k_* is a scalar weighting factor associated with the design objective of product *k*. Different Pareto optimal solutions can be obtained by varying these weights. The blended formulation always provides Pareto optimal solutions as long as *a_k_* > 0 (∀*k* ∈ *K*). In practice, the product priority, *a_k_*, can be determined by criteria such as product market value or “pathway readiness level” (i.e., certain pathways are easier to engineer than others).

#### 2.1.9 Goal attainment formulation

In the goal attainment problem,^23^ a target value is defined for each objective:

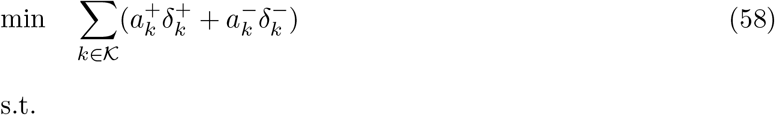

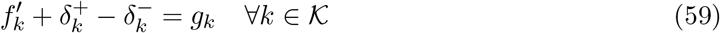

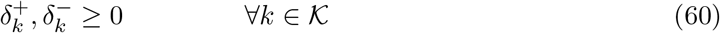

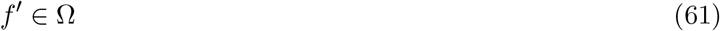

The problem seeks to minimize the variables 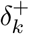 and 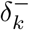 that represent the deficiency and excess of the objective 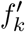 from the target value *g_k_*, respectively. Weighting parameters 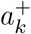 and 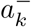 correspond to different types of discrepancy to be minimized. In general, when it is important to meet the target value without exceeding it, we set 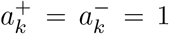; however, when the design objective is required to be greater or equal than the target value, we set 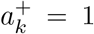 and 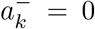, effectively converting (59) into 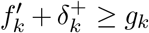. Solutions to the goal attainment problem are not guaranteed to be Pareto optimal, even if all demands *g_k_* are met. To address this issue, the blended problem (57) can be solved where the objectives are constrained to be equal or greater than the values found by solving the goal attainment problem. In practice, the goal attainment formulation corresponds to the identification of the modular cell *compatible* with the largest number of modules. Here, a module *k* is said to be *compatible* if 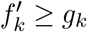.

### 2.2 Implementation

#### 2.2.1 Metabolic models

We used two parent models from which production networks were built, including: i) a core metabolic model of *E. coli*^17^ to develop the ModCell2-MILP algorithm and compare with previous ModCell2 results,^13^ and ii) the iML1515 genome-scale metabolic model of *E. coli*^24^ for biosynthesis of a library of endogenous and heterologous metabolites, including 4 organic acids, 6 alcohols, and 10 esters (Table 2).^25–34^ These models were configured as in the previous ModCell2 study^13^, briefly: Anaerobic conditions were imposed by setting oxygen exchange fluxes to be 0, and the glucose uptake rate was constrained to be at most 10 mmol/gCDW/h. When using the genome-scale model iML1515 to simulate *wGCP* designs, only the commonly observed fermentative products (acetate, CO_2_, ethanol, formate, lactate, succinate) were allowed for secretion as described elsewhere.^35^

#### 2.2.2 ModCell2-MILP simulations

ModCell2-MILP was implemented using Pyomo,^36^ an algebraic modeling language embedded in the Python programming language. All simulations were performed on a computer with an Intel Core i7-3770 processor, 32 GB of random access memory, and the Arch Linux operative system. The implementation and scripts used to generate the results of this manuscript are available as part of the ModCell2 package via Supplementary Material 2 and https://github.com/trinhlab/modcell2.

#### 2.2.3 Optimization solver configuration

The Pyomo^36^ implementation of ModCell2-MILP was solved with IBM Ilog Cplex 12.8.0. To avoid incorrect solutions associated with numerical issues the following Cplex parameters were changed from their default values: (i) *numerical emphasis* was set to “true”, (ii) *integrality tolerance* was lowered to 10^-7^, and (iii) the *MIP pool relative gap* was increased to 10^-4^ for enumerating alternative solutions. Alternative solutions were enumerated using the Cplex “populate” procedure.

### 2.3 Analysis methods

#### 2.3.1 Reference flux distribution

The reference flux distribution, 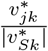, is determined by solving the following quadratic program based on the parsimonious enzyme usage hypothesis:^37,38^

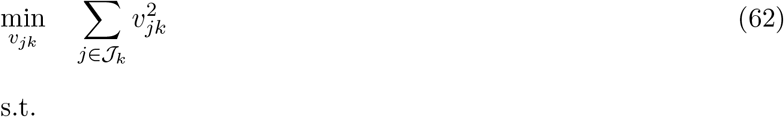

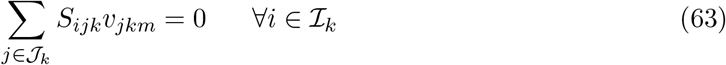

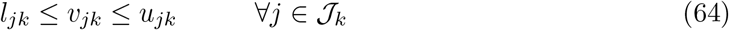

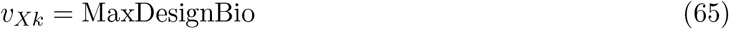

Constraint (63) corresponds to mass balance for the metabolic network. Constraint (64) corresponds to reaction bounds, including reaction deletions found in the modular cell design problem. Constraint (65) fixes the biomass formation rate, *v_Xk_*, to the maximum reachable by the design. This value (MaxDesignBio) is obtained by maximizing *v_Xk_* subject to (63) and (64). The reference flux distribution 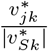 represents the desired metabolic state of a *wGCP* designed production network. This distribution, if feasible, is unique because the convex optimization problem is formulated with a positive definite quadratic objective function (see Theorem 16.4 in Nocedal and Wright^39^).

#### 2.3.2 Flux sampling

To determine an ensemble of flux distributions for a production network, we used the ACHR algorithm^40^ in the COBRA toolbox.^41^ Constraints for flux sampling simulation include the reaction deletions and module reactions found in the ModCell design problem solution, a fixed substrate uptake rate of −10 mmol glucose/gCDW/hr, and a minimum product synthesis flux of 50% of its maximum value.

#### 2.3.3 Metabolic map drawing

Drawings of metabolic map were performed using the Escher^42^ tool (https://escher.github.io) that produces *svg* files. Coloring, highlighting candidate reactions, and other systematic adjustments of metabolic maps were done with the Python-based *lxml* module. Additional editing for visual enhancement was done with the Inkscape software.

## 3 Results

### 3.1 Performance and solution time optimization of ModCell2-MILP

#### 3.1.1 ModCell2-MILP can not only reproduce the results of the original ModCell2 formulation but also find more alternative solutions

To evaluate ModCell2-MILP, we compared its performance with the previously developed ModCell2 platform^13^ that solves the optimization problem with multi-objective evolutionary algorithms (MOEAs). As a basis of comparison, we used the same *E. coli* core metabolic model, maximum number of deletion reactions *α*, and maximum number of module-specific reactions *β*_k_ for both Mod-Cell2 and ModCell2-MILP. Due to fundamental differences in problem formulations for MOEA and MILP, we used the *lsGCP* design objective for ModCell2-MILP with multiple weighting factors, ak, specifically selected to reproduce previous results, in the blended formulation and the *sGCP* design objective for ModCell2 (Supplementary Material 1). The results showed that ModCell2-MILP could generate the same Pareto optimal designs like ModCell2. In addition, ModCell2-MILP enumerated a larger number of alternative solutions than ModCell2. For example, the design named *sGCP-5-0-6* generated by ModCell2 had 3 alternative solutions while ModCell2-MILP found 8 alternative solutions. By increasing *α* to 8 and *β* to 2, we could identify a utopia design (i.e., one solution with the maximum value for all objectives) with 192 alternative solutions, which significantly expands the possibilities for experimental implementation.

#### 3.1.2 Tuning MILP formulations significantly improves solution times

We considered three techniques that can improve solution times of ModCell2-MILP, including:

i. *Fixing the network feasibility indicator w_k_*. If all modules are expected to be compatible with a final ModCell design (i.e., 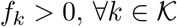), wk is set to be 1 for all 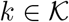 in order to avoid computational efforts in finding non-optimal feasible solutions.
ii. *Flux bound tightening*. Constraints of the form *e_jkm_l_jkm_* ≤ *v_jkm_* ≤ *e_jkm_u_jkm_* are known to result in weak linear relaxations, i.e., feasible values of *v_jkm_* are far from their bounds *l_jkm_* and *u_jkm_*. To tighten the formulation by making continuous relaxations closer to the feasible integer solution, smaller values of *u_jkm_* and *l_jkm_* are determined by solving a series of linear programs that maximize and minimize each flux *v_jkm_* in the parent production networks 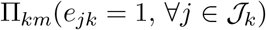.
iii. *Benders decomposition*. ModCell2-MILP has a separable structure compatible with Benders decomposition^43,44^ that creates a master problem, using binary variables and associated constraints (15)-(21), and sub-problems for each production network Φ*_km_*(*e_jk_*) with fixed binary variables. This decomposition implementation is automatically done by Cplex 12.8.

We evaluated these three techniques for tuning MILP formulations and used the core *E. coli* model^13^ for the benchmark study. The results showed that flux bound tightening, fixed *w_k_*, and Benders decomposition could reduce the solution time to find solutions by 50%, 80%, and 95%, respectively (Table 1). By combining these techniques, the solution time was shortened by 96% from 63.3 s to 2.8 s. In subsequent studies, we used these three tuning techniques to solve the ModCell design problem unless otherwise noted.

**Table 1:**
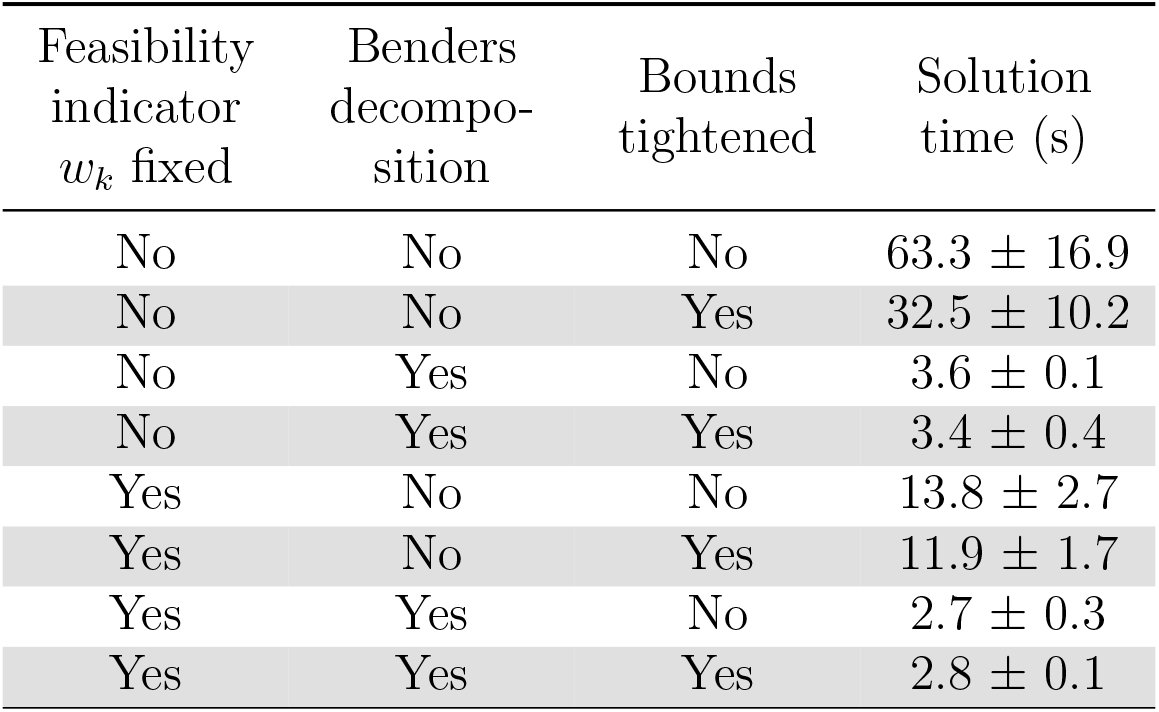
Solution time reduction by tuning the ModCell2-MILP formulation with Benders decomposition, bound tightening, and/or fixed network indicator 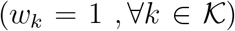. The simulations were performed in triplicates.

| Feasibility<br>indicator<br>$w_k$ fixed | Benders<br>decompo-<br>sition | Bounds<br>tightened | Solution<br>time (s) |
| --- | --- | --- | --- |
| No | No | No | $63.3 \pm 16.9$ |
| No | No | Yes | $32.5 \pm 10.2$ |
| No | Yes | No | $3.6 \pm 0.1$ |
| No | Yes | Yes | $3.4 \pm 0.4$ |
| Yes | No | No | $13.8 \pm 2.7$ |
| Yes | No | Yes | $11.9 \pm 1.7$ |
| Yes | Yes | No | $2.7 \pm 0.3$ |
| Yes | Yes | Yes | $2.8 \pm 0.1$ |

#### 3.1.3 Choices of design parameters affect solution time

In designing a modular cell with ModCell2-MILP, the designer needs to specify the formulation type (i.e., blended or goal attainment formulation), the target phenotype (e.g., *wGCP, lsGCP*, and *NGP*), and the limits of deletion reactions (α) and endogenous module-specific reactions *(β_k_*). We evaluated the impact of these parameters on solution time using the *E. coli* core model (Figure 2). Regardless of the formulation type, increasing α and β led to harder problems and hence required more solution time due to the exponentially increasing number of feasible solutions as expected. The goal attainment formulation took longer time to solve for the *lsGCP* and *NGP* design objectives, but about the same time for the *wGCP* design objective. Interestingly, the overall difficulty of *wGCP* is higher than that of *lsGCP* in both the blended and goal attainment formulations, despite *lsGCP* having approximately twice the number of constraints. Furthermore, the *NGP* design objective could be solved most quickly, likely due to the narrower design space associated with the no-growth associated production of target metabolites.

**Figure 2:**
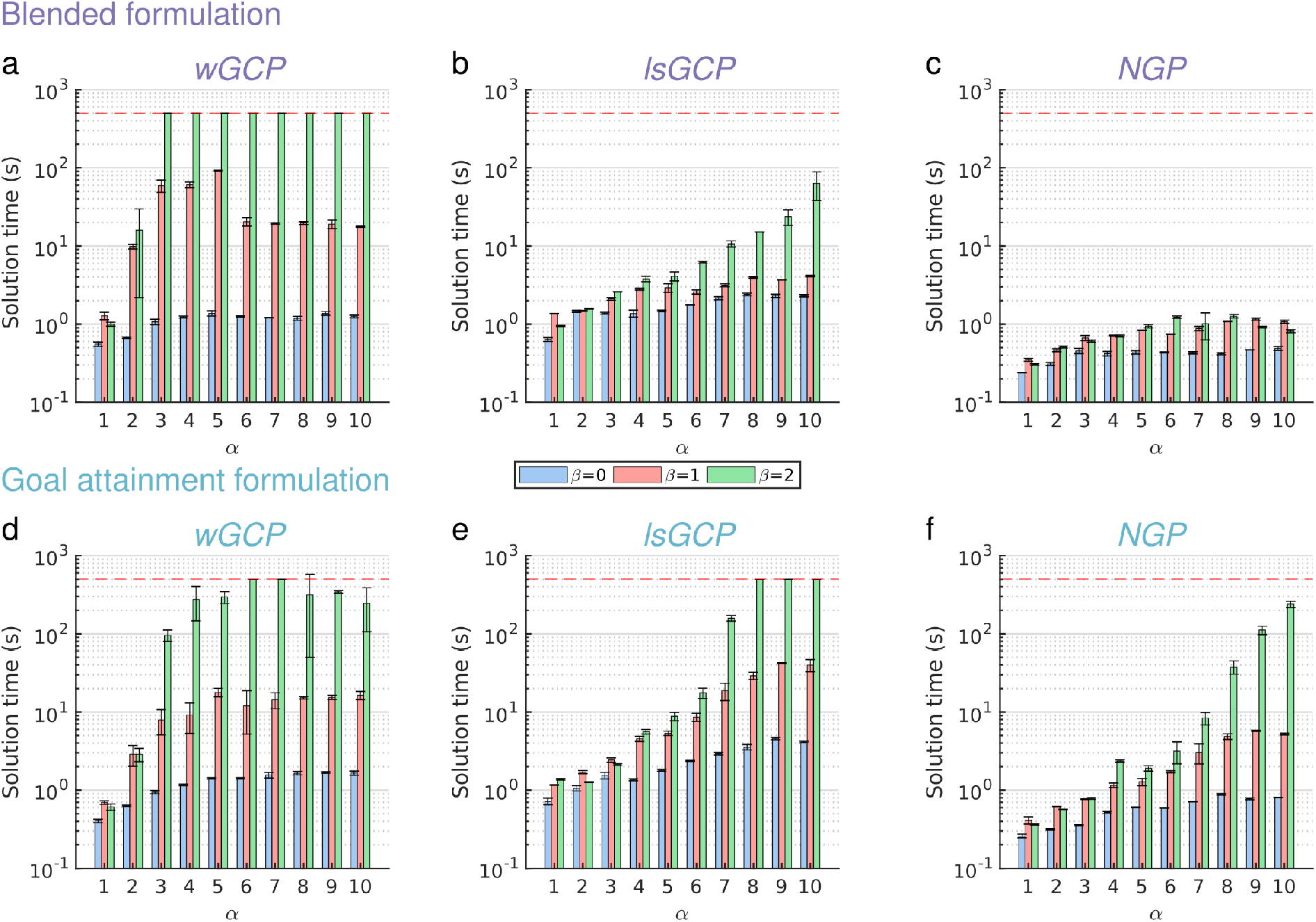
Effect of design parameters, including the target design objective (i.e., *wGCP, lsGCP*, and *NGP*) and the limits of deletion reactions *α* and endogenous module-specific reactions *β_k_*, on computation time for solving the ModCell2-MILP problem with the blended (a-c) and goal attainment (d-f) formulations. A time limit of 500 seconds indicated by a red dashed line was used in all cases, but only reached by certain *wGCP* and *lsGCP* cases with *β* ≥ 2. The simulations were performed in duplicates.

### 3.2 Design of a universal modular cell for a genome-scale metabolic model of *E. coli*

#### 3.2.1 Reduction of the candidate reaction deletion set enables ModCell2-MILP to find modular cell designs for a large-scale metabolic network

Finding genetic modifications towards a desired phenotype using mathematical optimization for large-scale metabolic networks has been known to be a computationally expensive task due to the combinatorial search space spanned by a large number of reaction deletion candidates in the network.^21,45^ Preprocessing of metabolic networks to reduce reaction candidates is not only critical but also practical for experimental implementation. Previous implementation of ModCell2 for the latest genome-scale *E. coli* model (iML1515)^24^ showed that the preprocessing step could reduce the set of reaction candidates from 2,712 to 276. By using ModCell2 with the *wGCP* objective, an *E. coli* modular cell was identified to be compatible with 17 out of 20 products with requirement of only 4 reaction deletions.^13^ Since MOEA implemented in ModCell2 does not guarantee optimality, here we aimed to evaluate the capability of ModCell2-MILP for handling a large-scale metabolic network and identifying the Pareto optimality and potential alternative solutions.

We applied ModCell2-MILP to analyze the same iML1515 model with a set of 20 products using the same design parameters (i.e., *α* and *β_k_*) and the blended formulation with all objective weights *α_k_* = 1. The simulation shows that ModCell2-MILP could not solve the ModCell design problem to optimality over 2 days of run time, likely due to the large number of candidate deletion reactions still present in the genome-scale model. To address this problem, the set of candidate reactions must be further reduced. Since only a small subset of all metabolic reactions in genome-scale models tend to be deleted by strain design algorithms,^13,21,46^ we used a pool of *wGCP* designs with *α* = 4, 5,6 and *β* = 0, 1 reported with ModCell2^13^ to identify relevant deletion candidates. From a set of 601 designs found by ModCell2, only 33 out of 276 candidates reaction deletions were used at least once. Hence, these 33 reactions were used to create a new, computationally-tractable set of reaction candidates. This new set contains reactions mostly from the well-characterized central metabolic pathways (Figure 3a) while the original set includes reactions in peripheral pathways that lead to biomass synthesis. Interestingly, within these 33 reaction candidates, only a few are used in most designs (Figure 3b), highlighting the importance of their removal in growth-coupled production phenotypes. Reactions with high deletion frequencies mainly occur in high-flux central metabolic pathways (Figure 3c), closely associated with cellular energetics and carbon precursors that interface with the production modules (Figure 3d).

**Figure 3:**
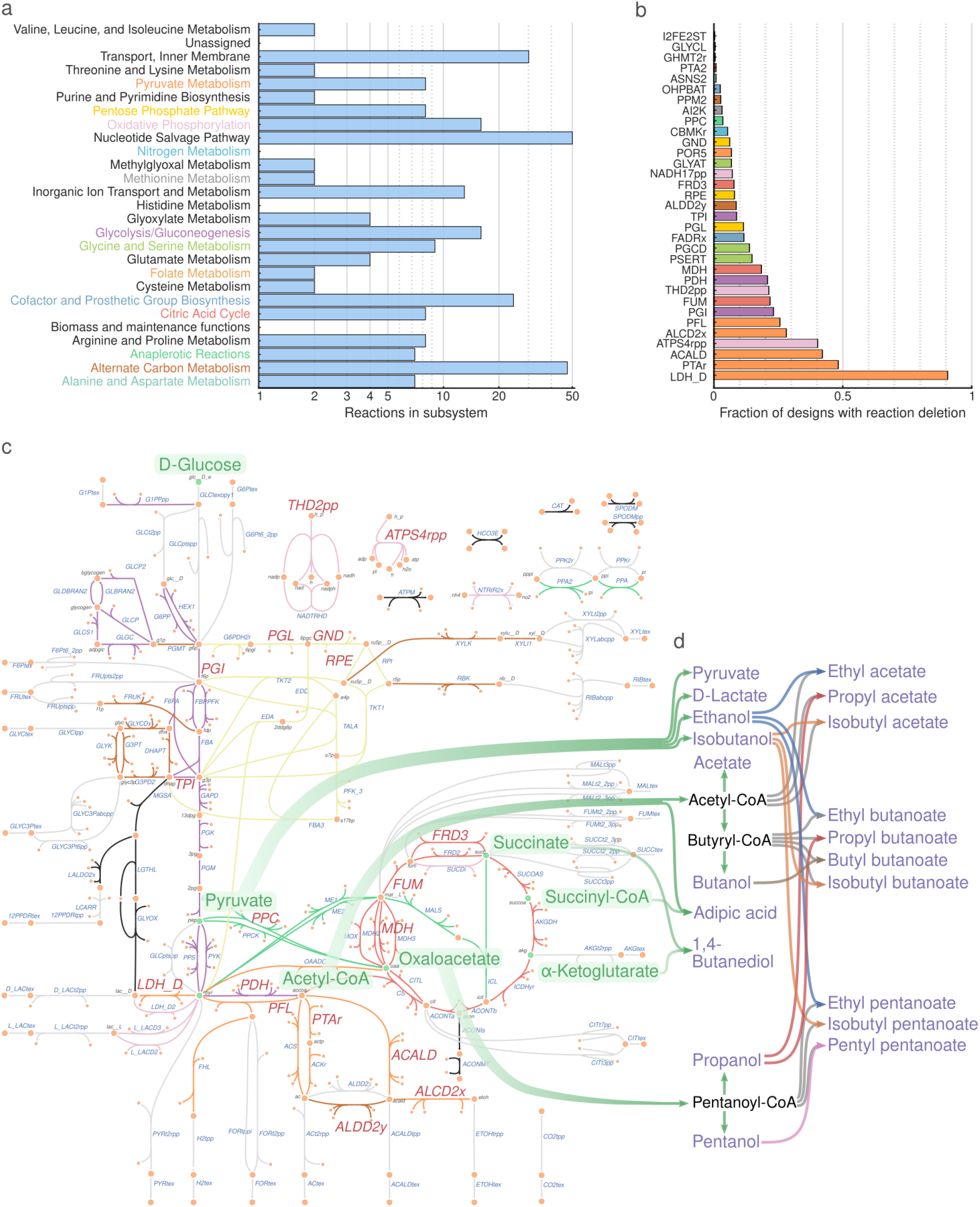
Analysis of reaction deletion candidates. (a) Subsystem distribution for the original set of 276 candidate reactions in the iML1515 model. Those subsystems that contain a reaction used in at least one design are colored. (b) Deletion frequency for the reduced set of 33 candidate reactions. The analysis is based on a pool of 601 *wGCP* designs from different *α* and *β* parameters whose Pareto fronts were previously determined with ModCell2.^13^ Bar colors indicate membership of these reactions to the subsystems. (c) Metabolic map of core metabolism. Key metabolites, including precursors for the 20 product modules (i.e., pyruvate, acetyl-CoA, succinyl-CoA, succinate, and *α*-ketoglutarate), are highlighted in green. Reactions are colored according to subsytem labels indicated in (a), reactions colored in light gray do not appear in any of the subsytems of (a), and reactions that are candidates for deletion, listed in (b), are labeled in red. (d) Link between major precursors and target products where colors are only used to facilitate visualization. Reaction and metabolite abbreviations correspond to BiGG^74^ identifiers (http://bigg.ucsd.edu/).

Using the reduced candidate reaction deletion set, ModCell2-MILP could find an optimal solution in ~ 30 min and enumerated all optimal solutions in ~ 8 hours. All the optimal solutions found by ModCell2-MILP in this case were in agreement with those previously found in ModCell2.^13^

#### 3.2.2 ModCell2-MILP can identify a universal modular cell compatible with all exchangeable production modules

Based on the computationally-tractable candidate reaction deletion set, we next evaluated whether the goal programming formulation could help identify a universal ModCell design that is compatible with all modules. By screening for various *α* and *β_k_*, we identified a universal modular cell that is compatible with all production networks, corresponding to the defined minimum design objective goal of 0.5 (i.e., 50% of the theoretical maximum product yield attained at the maximum growth rate), *α* = 6, and *β* = 1 (Figure 4a). Remarkably, most products greatly overcame this minimum goal with yields above 90% of the theoretical maximum values (Figure 4b). All production networks displayed a feasible metabolic space where an increase in product synthesis rate is needed to attain faster growth rates (Figure 4c). This designed phenotype is useful for optimal pathway selection using adaptive laboratory evolution^47,48^ and/or pathway libraries.^49^

**Figure 4:**
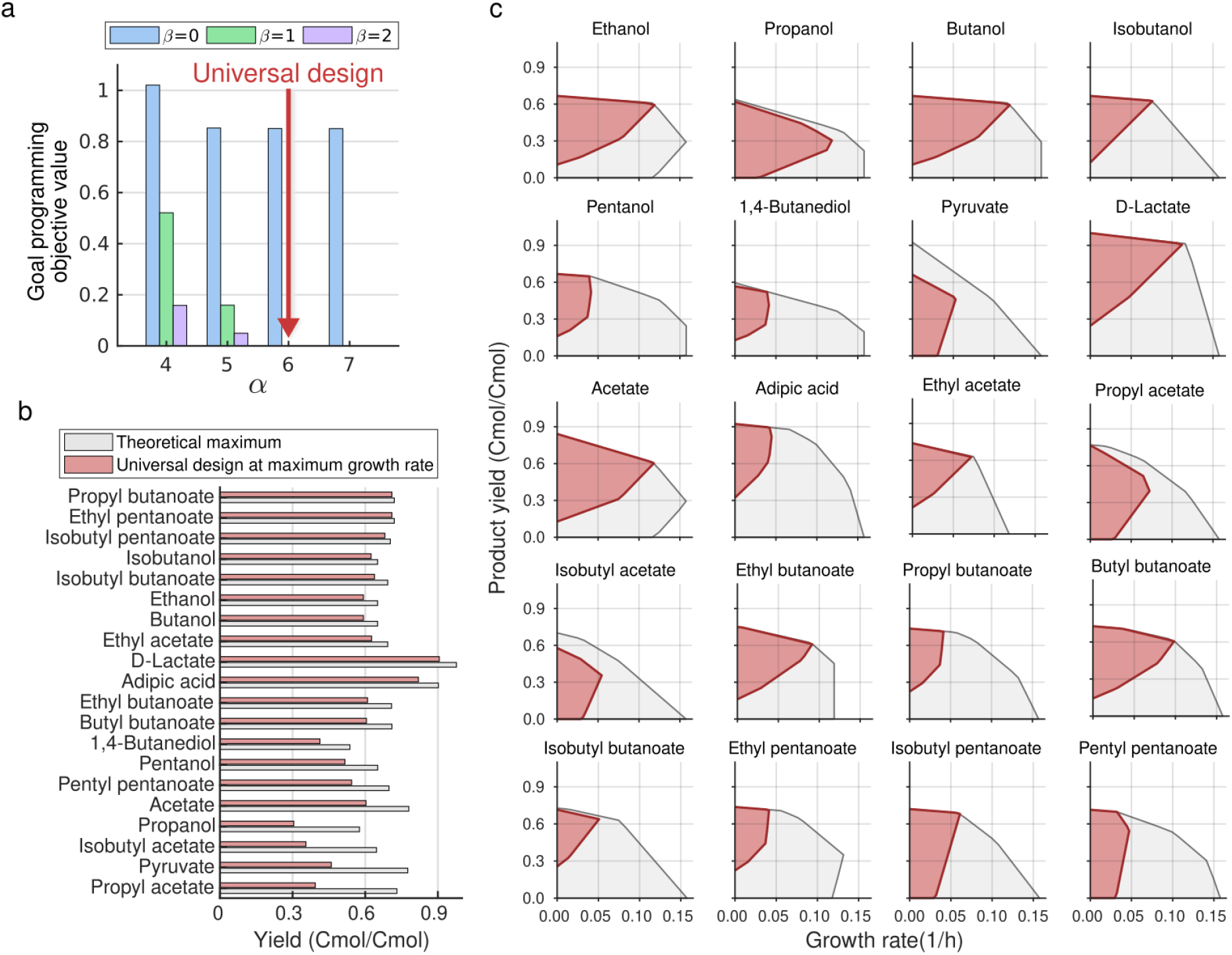
Identification of a universal modular cell compatible with all production modules using the *wGCP* design objective. (a) Goal programming solutions with increasing *α* and *β* values. The goal programming objective value (58) in the y-axis measures the difference between the performance of production strains and the target goal, i.e., 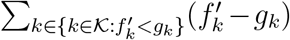 where the target goal is set to be *g_k_* = 0.5. The parameters *α* = 6 and *β* = 1 are sufficient to identify a universal modular cell design meeting the required goal for all production networks. (b) Comparison between the yield performances of the designed modular production strains and maximum theoretical values. (c) The feasible flux spaces for the wild-type (gray) and designed modular production strains (crimson). Based on the *wGCP* design phenotype, to increase growth rate, each mutant must increase product synthesis rate. The genetic manipulations of this universal modular cell design are indicated in the metabolic map of Figure 5c.

### 3.3 Flexible metabolic flux capacity of *E. coli* core metabolism enables the design of a universal modular cell

#### 3.3.1 Endogenous modules responsible for metabolic flexibility of a universal modular cell are identified by comparing flux distributions of production networks

The designed universal modular cell (Section 3.2.2) can theoretically adapt to the contrasting metabolic requirements of all production modules (Table 2). To gain further insight into this unique metabolic capability of the modular cell and its potential to be realized in practice, we analyzed its *reference flux distributions* (Section 2.3.1) across the production networks. Reactions with the highest flux changes across the production networks are likely critical for the proper operation of the universal modular cell and might present potential bottlenecks. Such reactions were identified by filtering their reference flux standard deviation (calculated across production networks) with an *ad hoc* threshold of 0.2 (mol/substrate mol). Over 90% of the 535 active reactions, each of which carries a non-zero flux in at least one production network, had standard deviation values below the threshold, indicating highly conserved metabolic core pathways among production networks. Only 9.5% of the active reactions presented a standard deviation magnitude above the threshold (Figure 5a).

**Table 2:** Overall production module stoichiometries, degree of reduction (DoR) of the final product (mol *e*^-^ / mol C), and metabolite secretion profiles from simulated reference flux distributions (mol C / mol C) of the universal modular cell design. Flux (mmol/gCDW/hr) abbreviations: *r_p_*, product; *r_ac_*, acetate; *r*_*co*_2__, CO_2_; *r_for_*, formate; *r_succ_*, succinate. Note that the negative CO_2_ fluxes in pyruvate and acetate production networks indicate overall CO_2_ uptake enabled by phosphoenolpyruvate carboxylase (PPC).

| Overall reaction | DoR | $r_p$ | $r_{ac}$ | $r_{co_2}$ | $r_{for}$ | $r_{succ}$ |
| --- | --- | --- | --- | --- | --- | --- |
| pyr + nadh → **ethanol** \| accoa + 2 nadh → **ethanol** (native) | 7.0 | 0.58 | 0.01 | 0.27 | 0.04 | - |
| oaa + glu + 2 atp + 2 nadph + nadh → akg + **propanol** | 6.7 | 0.31 | 0.36 | 0.07 | 0.18 | - |
| 2 accoa + 4 nadh → **butanol** | 6.5 | 0.59 | 0.01 | 0.28 | 0.04 | - |
| 2 pyr + nadph + nadh → **isobutanol** | 6.5 | 0.62 | - | 0.31 | - | - |
| oaa + glu + accoa + 3 nadh + 2 atp + 2 nadph → akg + **pentanol** | 6.4 | 0.50 | 0.21 | 0.24 | 0.03 | - |
| succ + akg + atp + 4 nadh + accoa → ac + **1,4-butanediol** | 5.5 | 0.46 | 0.33 | 0.17 | - | - |
| → **pyruvate** | 3.0 | 0.46 | - | -0.16 | - | 0.66 |
| pyr + nadh → **D-lactate** | 3.7 | 0.91 | - | - | - | - |
| accoa → atp + **acetate** | 3.5 | 0.60 | 0.60 | -0.30 | 0.61 | - |
| accoa + succoa + 2 nadh → atp + **adipic acid** | 4.0 | 0.82 | 0.05 | 0.04 | 0.06 | - |
| accoa + pyr + nadh → **ethyl acetate** | 5.0 | 0.63 | - | - | 0.32 | - |
| accoa + oaa + glu + 2 atp + 2 nadph + nadh → akg + **propyl acetate** | 5.2 | 0.41 | 0.30 | - | 0.24 | - |
| accoa + 2 pyr + nadph + nadh → **isobutyl acetate** | 5.3 | 0.36 | - | 0.02 | 0.06 | 0.52 |
| 2 accoa + 3 nadh + pyr → **ethyl butanoate** | 5.3 | 0.61 | - | 0.09 | 0.23 | - |
| 2 accoa + 3 nadh + oaa + glu + 2 atp + 2 nadph → akg + **propyl butanoate** | 5.4 | 0.68 | 0.03 | 0.23 | 0.04 | - |
| 4 accoa + 6 nadh → **butyl butanoate** | 5.5 | 0.61 | - | 0.14 | 0.18 | - |
| 2 accoa + 3 nadh + 2 pyr + nadph → **isobutyl butanoate** | 5.5 | 0.64 | - | 0.16 | 0.16 | - |
| oaa + glu + accoa + 2 nadh + 2 atp + 2 nadph + pyr → akg + **ethyl pentanoate** | 5.4 | 0.68 | 0.03 | 0.23 | 0.04 | - |
| oaa + glu + accoa + 2 nadh + 2 atp + 3 nadph + 2 pyr → akg + **isobutyl pentanoate** | 5.6 | 0.67 | 0.01 | 0.25 | 0.03 | - |
| 2 oaa + 2 glu + 2 accoa + 4 nadh + 4 atp + 4 nadph → 2 akg + **pentyl pentanoate** | 5.6 | 0.53 | 0.22 | 0.20 | 0.02 | - |

**Figure 5:**
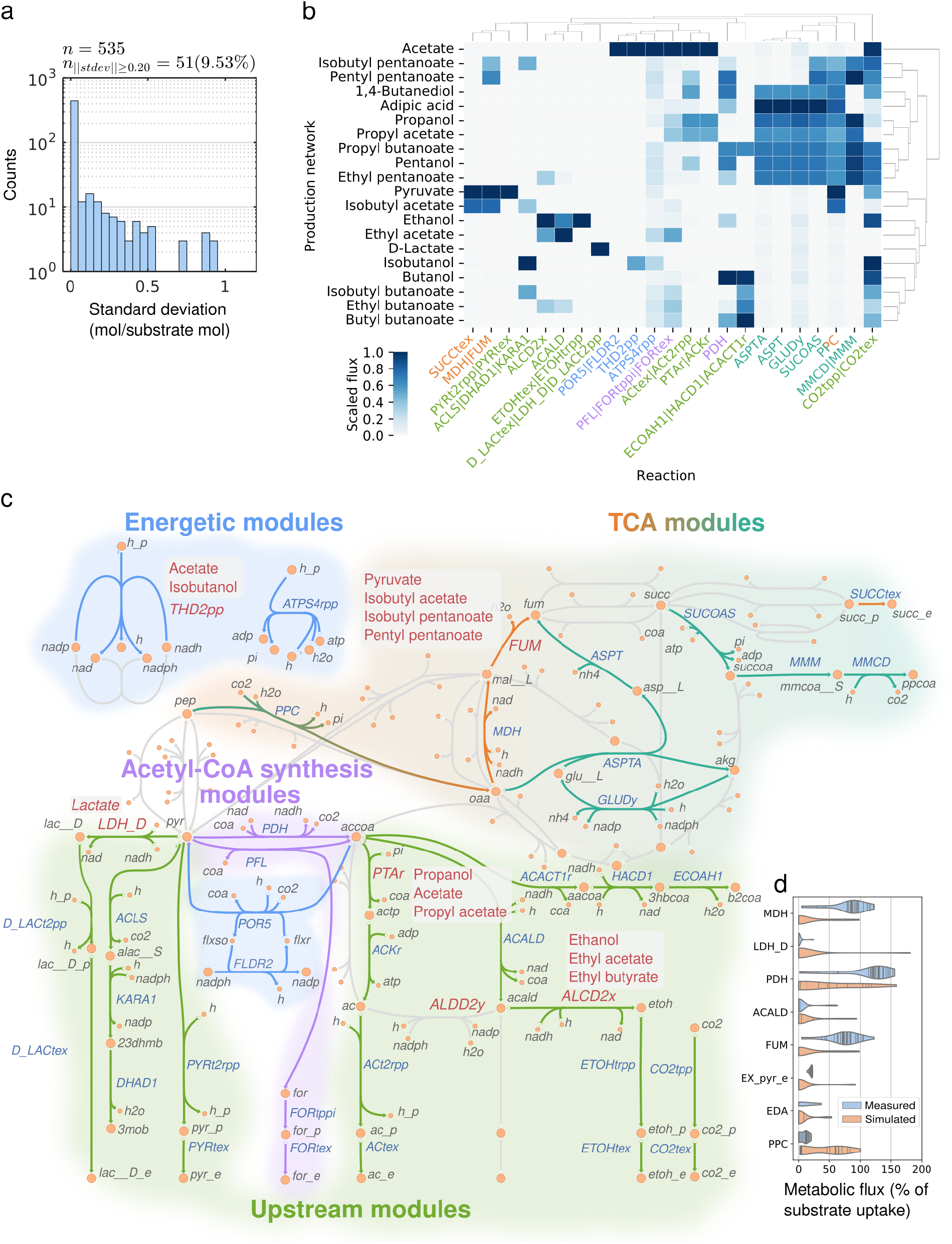
Flexible metabolic flux capacity of *E. coli* metabolism enables the universal modular cell design. (a) Standard deviation of each reaction flux across production networks. (b) Scaled fluxes of the 51 reactions with standard deviation magnitude above 0.2, excluding proton, water transport, and exchange reactions. A scaled flux for a reaction is determined by the reference flux distribution value divided by the maximum value of that reaction across all production networks. Thus, a scaled flux of 0 indicates a given reaction does not carry any flux, and a scaled flux of 1 indicates that this reaction carries the highest flux across production networks. Several columns have multiple reactions, separated by |, since they carry exactly the same flux. (c) Endogenous modules of the universal modular cell. The reactions colored in red are deleted in the chassis, but are used as module reactions in the production networks shown in the adjacent gray boxes. Metabolites in periplasmic and extracellular compartments have “_p” and “_e” suffixed to their abbreviations, respectively. Metabolite and reaction abbreviations follow BiGG^74^ notation. (d) Comparison between simulated and measured fluxes. The solid lines within the “violins” correspond to samples. The simulated fluxes for the reversible reactions, including FUM, LDH, MDH, and ACALD, were multiplied by −1 to reflect their most common direction.

In our case study of designing a universal modular cell compatible with all 20 production modules, unbiased clustering analysis (Figure 5b) revealed the presence of four endogenous module types in the core metabolism of *E. coli* that are activated to fit specific production modules (Figure 5c). In the context of chassis metabolism, an endogenous module corresponds to a reaction or group of highly coupled reactions that become active to accomplish a certain metabolic function. The endogenous module classification can be understood in terms of location (i.e., proximity in the metabolic network) and three metabolic functions. The first function is the direction of carbon towards general precursor metabolites including (i) pyruvate and acetyl-CoA captured by acetyl-CoA-associated modules and (ii) oxaloacetate, succinate, succinyl-CoA, and *α*-ketoglutarate captured by TCA-associated modules. The second function is the direction of carbon from the precursor metabolites towards secretable molecules, captured by the upstream and TCA-associated modules. The third function is the use of ATP- and NADP(H)-dependent pathways required to maintain homeostasis, captured by the acetyl-CoA-associated and energetic modules. While these functions are conceptually separable, their biochemical manifestation overlaps, i.e., specific metabolic reactions or pathways can simultaneously fulfill several functions.

Each endogenous module can be viewed as an interface of the universal modular cell with production modules that are exchangeable. The endogenous modules might become potential metabolic bottlenecks in practice if they cannot satisfy the predicted fluxes, and thus might be critical engineering targets when the associated production modules are used.

##### Acetyl-CoA-associated endogenous modules

This module type contains pyruvate formate lyase (PFL) and pyruvate dehydrogenase enzyme complex (PDH) reactions that convert pyruvate to acetyl-CoA. Intuitively, products derived from pyruvate, such as isobutanol, require a low flux through PFL and PDH while those derived from acetyl-CoA require a high flux. Remarkably, the redox states of production strains determine the ratios of PFL to PDH fluxes. For example, the ethanol production network has a relatively high flux through PDH and a low flux through PFL; however, for ethyl acetate that has a lower degree of reduction than ethanol (Table 2), PFL with formate secretion is prioritized over PDH with NADH generation. Note that our model did not include the regulatory restriction that PDH is inhibited in *E. coli* anaerobically because the function of PDH is equivalent with the coupling of PFL and heterologous NADH-dependent formate dehydrogenase (FDH) demonstrated experimentally for increased butanol^30,50^ and pentanol^32^ production.

##### Upstream modules

This module type is formed by reactions located directly upstream of a secretable metabolite, often associated with the target production module, and thus provides the necessary precursor metabolite(s). Such reactions are commonly over-expressed in practice, e.g., the ECOAH1-HACD1-ACACT1r endogenous module (comprising of 3-hydroxyacyl-CoA dehydratase, 3-hydroxyacyl-CoA dehydrogenase, and acetyl-CoA acetyl transferase) responsible for generating butyryl-CoA and the ACLS-DHAD1-KARA1 endogenous module (comprising of acetolactate synthase, dihydroxy-acid dehydratase, and keto-acid reductoisomease) responsible for generating isobutyryl-CoA. These endogenous modules can also become active to form byproducts in certain production networks, e.g., the PTAr-ACKr-ACT2rpp-ACtex endogenous module (comprising of phosphate acetyl transferase, acetate kinase, and cytosolic and periplasmic acetate transport) that not only carries the highest flux in the acetate production network but also becomes active in the propanol-associated modules.

##### TCA-associated endogenous modules

This module type has the same function as the upstream endogenous modules but it is localized in the TCA (Krebs) cycle. Several products, including adipic acid, 1,4-butanediol, propanol, pentanol, and their associated esters, are derived from the TCA intermediates and interface with the universal modular cell via the TCA-derived endogenous modules. The SUCOAS-MMM-MMCD endogenous module (comprising of succinyl-CoA synthetase, Methylmalonyl-CoA mutase, methylmalonyl-CoA decarboxylase) must be activated to convert succinate into succinyl-CoA and then propanoyl-CoA. Remarkably, two routes are present to synthesize fumarate from oxaloacetate, including the conventional MDH-FUM endogenous module (comprising of malate dehydrogenase and fumarase) that consumes NADH and the cyclic ASPTA-GLUDY-ASPT endogenous module (comprising of aspartate transaminase, glutamate dehydrogenase, and L-aspartase) that consumes NADPH. These NADH/NADPH cofactors are not interchangeable due to the deletion of the transhydrogenase THD2pp in the universal modular cell, so the isobutyl pentanoate and pentyl pentanoate modules, that are derived from the ASPTA-GLUDY-ASPT endogenous module, also have a high NADPH requirement. Some production networks, such as pyruvate and isobutyl acetate that are not based on the TCA-derived endogenous modules, secrete succinate instead of ethanol and/or lactate to balance redox by using the PPC-MDH-FUM-SUCCtex endogenous module (comprising of phosphoenolpyruvate carboxylase, malate dehydrogenase, fumarase, and succinate transport).

##### Energetic modules

This module type primarily involves NAD(P)-dependent transhydrogenase (THD2pp) and ATP synthase (ATPS4rpp). Other reactions that allow coupling of phosphate- and electron-transfer cofactors are also included. The reactions in this module help buffer the diverse electron and ATP requirements of production networks. THD2pp is deleted in the chassis but used as a module reaction in the isobutanol and acetate production networks. In the case of isobutanol production, transhydrogenase expression has been demonstrated to increase the synthesis of NADPH and thus isobutanol.^51^ Acetate has the smallest degree of reduction after pyruvate, which results in redox imbalance that is compensated via formate secretion. In conjunction with these mechanisms, ATPsynthase works in the reverse direction by hydrolyzing excess ATP. Other production networks also use ATPS4rpp to eliminate excess ATP as observed, for example, in the ethyl acetate production network. This strategy is consistent with ATP wasting approaches recently demonstrated.^52^

#### 3.3.2 Comparison between simulated and measured intracellular fluxes reveals flexible metabolic flux capacity of *E. coli* to accommodate the required wide flux ranges

Flux analysis of the production networks suggests that the core metabolic reactions (Figure 5b) require a wide range of fluxes. To successfully implement this modular design in practice, we need to evaluate whether the metabolism of *E. coli* has the inherent metabolic flux capacity to accommodate the required fluxes of the designed universal modular cell when coupled with various exchangeable production modules. We compared the simulated reference flux distributions with a recent collection of 45 measured metabolic fluxes^53^ that are collected from multiple studies across various conditions (e.g., growth under aerobic and anaerobic conditions, use of glucose or acetate or pyruvate as a carbon source) and genotypes (e.g., wild-type *E. coli* and mutants with single gene deletions).^54–57^ Note that this dataset provides a baseline for wild-type and relatively small deviations from that state (i.e., single gene deletion mutants), thus highly engineered strains (e.g., with three or more gene deletions) are likely to attain wider flux distributions.

Within the 23 reaction groups that constitute endogenous modules (Figure 5b), 8 reactions could be matched to this experimental dataset (Figure 5d). Remarkably, a highly consistent overlap of flux ranges was observed between the simulated and measured fluxes for malate dehydrogenase (MDH), pyruvate dehydrogenase (PDH), acetaldehyde dehydrogenase (ACALD), fumarase (FUM), and 2-dehydro-3-deoxy-phosphogluconate aldolase (EDA). For the cases of D-lactate dehydrogenase (LDH_D), and pyruvate secretion (EX_pyr_e) that are directly coupled with the biosynthesis of lactate and pyruvate, respectively, we observed the maximum simulated fluxes surpass the measured values, suggesting that further engineering of wild-type and single-gene deletion *E. coli* is needed to attain the requried fluxes. Indeed, previous studies^58,59^ have been able to redirect metabolic fluxes in *E. coli* for yields of lactate and pyruvate above 75% of the theoretical maximum values by simultaneous elimination of competing fermentative pathways, including acetate (Δ*ackA*), formate (Δ*pflB*), and ethanol (Δ*adhE*). The only remaining discrepancy between the simulated and measured fluxes is PPC. Studies, not included in the comparison data set, have reported up to 50% more PPC flux observed under aerobic conditions^60,61^, which is still considerably below several of the simulated fluxes. This result suggests that PPC can be a potential metabolic bottleneck in certain production modules. One potential solution is to include in the affected production modules the heterologous PPC from *Actinobacillus succinogenes* which has been successfully over-expressed in *E. coli* for increased succinate production.^62^ Additionally, bacterial PPC activity can be increased by elevating the acetyl-CoA pool.^63^

#### 3.3.3 Random sampling of metabolic fluxes confirms the narrow operation range of endogenous modules

The calculated reference flux distributions represent the ideal metabolic states for each production strain. However, other metabolic states might also exist. To address this uncertainty, we performed randomized flux sampling^40,41^ for each production network under the constraint that product synthesis rate has to be above 50% of the maximum value (Section 2.3.2). The results show that the metabolic flux distributions for most reactions involved in the endogenous modules (Figure 6a-u) are very narrow, except the two alternative pathways for ethanol synthesis, i.e., the endogenous PDH-ACALD-ALCD2x route (comprising of pyruvate dehydrogenase, acetaldehyde dehydrogenase, and alcohol dehydrogenase) (Figure 6t) route and the heterologous PDC-ALCD2x route (comprising of pyruvate decarboxylase and alcohol dehydrogenase). The range of experimental and simulated fluxes are comparable, which is consistent with the results in Section 3.3.2. In summary, even though reactions in the endogenous modules must have flexible metabolic flux capacities to enable a universal modular cell to be compatible with various exchangeable production modules, they must also operate within in a narrow flux range when interfacing with a specific production module.

**Figure 6:**
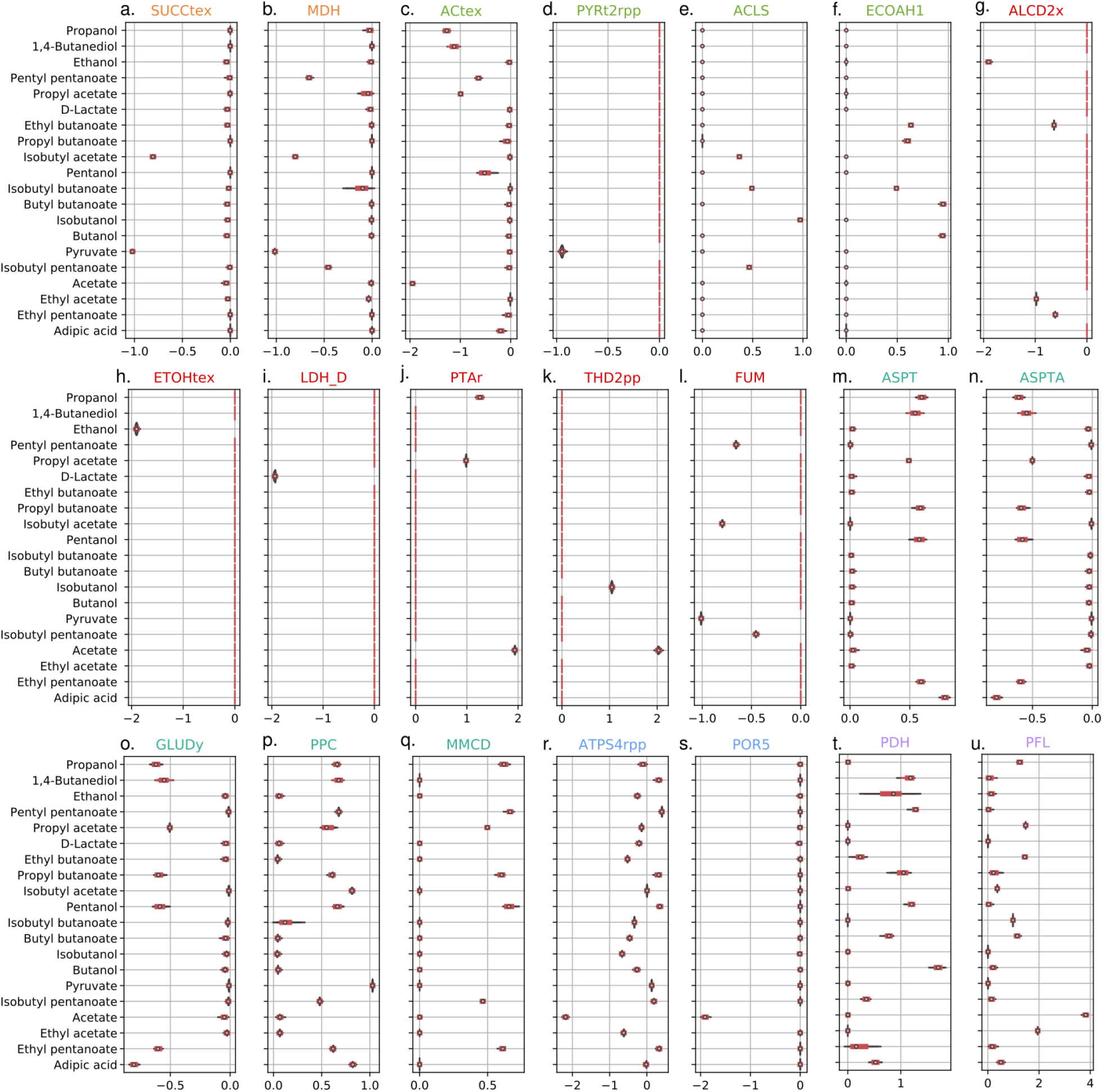
Violin plots of sampled flux distributions of the reactions of interest. Reaction colors are consistent with Figure 5. The flux of SUCOAS could not be sampled since this reaction is involved in a thermodynamically infeasible cycle.

## 4 Conclusions

Modular cell design seeks to accelerate strain development towards broader biotechnological application of synthetic biology and metabolic engineering, similar to the proven advantages of modular design in conventional engineering disciplines.^4^ In this study, we adapted the recently proposed^13^ multi-objective modular strain design method to a MILP computational framework that can guarantee Pareto optimal solutions, exhaustively search the space of alternative solutions, and specify design goals such as module prioritization. Remarkably, the proposed method identified a universal modular cell that harnesses the inherent modularity and flexibility of native *E. coli* metabolism^64,65^ to properly interface with a variety of biochemically diverse heterologous pathways. This universal design is predicted to display a growth-coupled to product formation phenotype for all pathways, enabling its use as a platform for pathway optimization through high-throughput library selection or adaptation. The feasibility of this universal design strategy is found to be consistent with experimental evidence of isolated metabolic engineering strategies towards target products and measured intracellular flux ranges. We anticipate this is the first example of upcoming methodological developments in the multi-objective strain design approach, which will follow a path similar to single-phenotype strain design algorithms^66^ introduced in the early 2000s,^18^ including the addition of heterologous metabolic reactions from large biochemical databases^67^ and up- and down-regulation of genes in addition to knock-outs^68^, as well as the use of alternative modeling paradigms for flux prediction such as kinetic models^69^ and ME-models.^70^ Additionally, we anticipate that the method developed in this study can be applied to exchangeable metabolic modules whose functions can be expanded to bioremediation^71^ and biosensing^72,73^.

## Acknowledgments

This research was funded by the NSF CAREER Award (NSF#1553250) and the Center of Bioenergy Innovation (CBI), U.S. Department of Energy Bioenergy Research Center supported by the Office of Biological and Environmental Research in the DOE Office of Science. The funders had no role in the study design, data collection and analysis, decision to publish, or preparation of the manuscript.

## 5 Definitions

### Sets

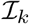 Metabolites in production network *k*.
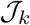 Reactions in production network *k*.
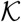 Production networks that are derived from a combination of the parent metabolic network with the metabolic pathways associated with production modules. The parent metabolic network is the network of the host strain that is genetically manipulated to build a modular cell chassis.
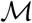 Metabolic states that correspond to the growth phase, denoted *μ*, and the non-growth or stationary phase, denoted 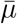.
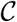 Candidate deletion reaction set. The removal of these reactions are applied to all production networks, 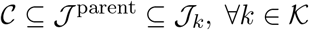.
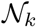 Non-targeted deletion reaction set in production network *k*. This set arises from the use of fixed endogenous module reactions *z_jk_* in certain production networks.

### Binary variables

*y_j_* Reaction deletion indicator that takes a value of 0 if reaction j is deleted in the chassis and 1 otherwise.
*z_jk_* Endogenous module reaction indicator that takes a value of 1 if reaction *j* is added back as module reaction in production network *k* and 0 otherwise.
*d_jk_* Reaction activity indicator that takes a value of 0 if reaction *j* in production network *k* might not carry a flux and 0 otherwise, thus *d_jk_* = *y_j_* ∨ *j*. This variable is declared as a continuous and linear constraints enforce the OR relation and thus makes the variable binary.
*w_k_* Production network feasibility indicator that takes a value of 0 if reaction deletions are ignored and the objective value is set to 0 for production network *k*, and a value of 1 otherwise.
*e_jk_* Reaction activity indicator adjusted to *w_k_* that takes the value of *j* if *w_k_* = 1 and a value of 1 if *w_k_* = 0, thus *e_jk_* = (*d_jk_* ∧ *w_k_*) ∨ ¬*w_k_*.
*r_jk_* Linearization variable, *r_jk_* = *d_jk_* ∨ *w_k_*.

### Continuous variables

*v_jkm_* Flux (mmol/gCDW/hr) of reaction *j* from network *k* at metabolic state *m*.
*v_Pkm_* Flux (mmol/gCDW/hr) of product synthesis reaction from network *k* at metabolic state *m*.
*v_Xkm_* Flux (mmol/gCDW/hr) of biomass synthesis reaction from network *k* at metabolic state *m*.
*f_k_* General objective function for production network *k* that can be represented by 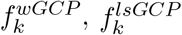, or 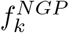.
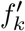 Objective function adjusted by *w_k_* such that 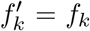 if *w_k_* = 1 and 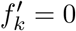 otherwise.
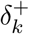 Amount required by the objective value 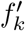 to attain the target goal *g_k_*, i.e.. 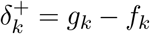 if 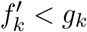.
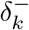 Amount that the objective value 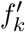 surpasses the target goal *g_k_*, i.e., 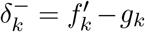 if 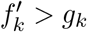.
*λ_ikm_* Dual variable associated with mass balance constraint of metabolite *i* from production network *k* at growth state *m*.
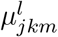 Dual variable associated with the lower bound of reaction *j* from production network *k* at growth state *m*.
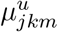 Dual variable associated with the upper bound of reaction *j* from production network *k* at growth state *m*.
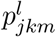 Linearization variable, 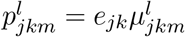.
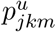 Linearization variable, 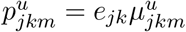.

### Parameters

*S_ijk_* Stoichiometric coefficient of metabolite *i* in reaction *j* of production network *k*.
*l_jkm_* Lower bound for reaction *j* of production network *k* at metabolic state *m*.
*u_jkm_* Upper bound for reaction *j* of production network *k* at metabolic state *m*.
*γ* Minimum biomass synthesis rate required for growth states. Note that in this study a conservative value of 20% of the maximum predicted growth rate of the wild-type strain was used to generate all results.
*α* Maximum number of deleted reactions in the modular cell chassis.
*β_k_* Maximum number of endogenous module reactions in production network *k*.
*ϵ* Small scalar used for tilting the biomass objective function, leading to the minimum product rate available at the maximum growth rate. Note that in our study *ϵ* = 0.0001 was used to generate all results.
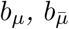 Weights on the growth and non-growth objectives of 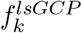, respectively. Note that in our study *b_μ_* = 1 and 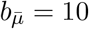 were used to generate all results.
*a_k_* Weighting factor applied to the objective function for production network *k* in the blended formulation. Note that in our study *a_k_* = 1, 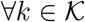 was used unless otherwise noted.
*g_k_* Target value for objective 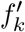 in the goal programming formulation.
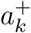 Weighting factor applied to 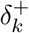 which emphasizes the importance of objective value 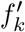 to avoid falling below the target value *g_k_*. Note that in our study 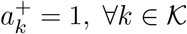 was used in all cases.
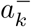 Weighting factor applied to 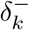 which emphasizes the importance of the objective 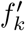 to avoid surpassing the target value *g_k_*. Note that in our study 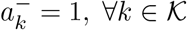 was chosen everywhere except to determine the universal modular cell design, where 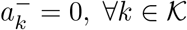 was used.
*M^w^* Determines the minimum value of *f_k_* that allows *w_k_* to not be 0. A value of 10, corresponding to *f_k_* > 0.01 for *w_k_* = 0, was used in all cases.
*M^obj^* Upper bound for each objective value. Note that in our study a value of 20 was set for all cases.
*M* Upper bound for dual variables. Note that in our study a value of 100 was set for all cases.

## 6 Reaction abbreviations

Identifier: Name
ACACT1r: Acetyl-CoA C-acetyltransferase
ACACT2rpp: Acetate reversible transport via proton symport (periplasm)
ACALD: Acetaldehyde dehydrogenase (acetylating)
ACKr: Acetate kinase
ACLS: Acetolactate synthase
ACtex: Acetate transport via diffusion (extracellular to periplasm)
ALCD2x: Alcohol dehydrogenase (ethanol)
ASPTA: Aspartate transaminase
ASPT: L-aspartase
ATPS4rpp: ATP synthase (four protons for one ATP) (periplasm)
DHAD1: Dihydroxy-acid dehydratase (2,3-dihydroxy-3-methylbutanoate)
ECOAH1: 3-hydroxyacyl-CoA dehydratase (3-hydroxybutanoyl-CoA)
EDA: 2-dehydro-3-deoxy-phosphogluconate aldolase
FUM: Fumarase
HACD1: 3-hydroxyacyl-CoA dehydrogenase (acetoacetyl-CoA)
KARA1: Ketol-acid reductoisomerase (2,3-dihydroxy-3-methylbutanoate)
MDH: Malate dehydrogenase
MMCD: Methylmalonyl-CoA decarboxylase
MMM: Methylmalonyl-CoA mutase
PDH: Pyruvate dehydrogenase
PFL: Pyruvate formate lyase
PPC: Phosphoenolpyruvate carboxylase
PTAr: Phosphotransacetylase
SUCCtex: Succinate transport via diffusion (extracellular to periplasm)
SUCOAS: Succinyl-CoA synthetase (ADP-forming)
THD2pp: NAD(P) transhydrogenase (periplasm)

## Supplementary Materials

**Supplementary Material 1** Modular cell designs for *E. coli* core model.

**Supplementary Material 2** Computer programs used to generate the results of this study.

## References

1. Lee, S. Y. et al. A comprehensive metabolic map for production of bio-based chemicals. Nature Catalysis 2, 18 (2019).

2. Nielsen, J. & Keasling, J. D. Engineering Cellular Metabolism. Cell 164, 1185–1197 (2016).

3. Trinh, C. T. & Mendoza, B. Modular cell design for rapid, efficient strain engineering toward industrialization of biology. Current Opinion in Chemical Engineering 14, 18–25 (2016).

4. Garcia, S. & Trinh, C. T. Modular design: Implementing proven engineering principles in biotechnology. Biotechnology Advances (2019).

5. Coello, C. A. C. & Lamont, G. B. Applications of multi-objective evolutionary algorithms (World Scientific, Singapore, 2004).

6. Rangaiah, G. P. Multi-objective optimization: techniques and applications in chemical engineering (World Scientific, Singapore, 2009).

7. Kitano, H. Biological robustness. Nature Reviews Genetics 5, 826 (2004).

8. Kashtan, N., Noor, E. & Alon, U. Varying environments can speed up evolution. Proceedings of the National Academy of Sciences 104, 13711–13716 (2007).

9. Clune, J., Mouret, J.-B. & Lipson, H. The evolutionary origins of modularity. Proc. R. Soc. B 280, 20122863 (2013).

10. Shoval, O. et al. Evolutionary trade-offs, Pareto optimality, and the geometry of phenotype space. Science, 1217405 (2012).

11. Schuetz, R., Zamboni, N., Zampieri, M., Heinemann, M. & Sauer, U. Multidimensional optimality of microbial metabolism. Science 336, 601–604 (2012).

12. Helmer, R., Yassine, A. & Meier, C. Systematic module and interface definition using component design structure matrix. Journal of Engineering Design 21, 647–675 (2010).

13. Garcia, S. & Trinh, C. T. Multiobjective strain design: A framework for modular cell engineering. Metabolic Engineering 51 (2019).

14. Garcia, S. & Trinh, C. T. Comparison of Multi-Objective Evolutionary Algorithms to Solve the Modular Cell Design Problem for Novel Biocatalysis. Processes 7 (2019).

15. Zhou, A. et al. Multiobjective evolutionary algorithms: A survey of the state of the art. Swarm and Evolutionary Computation 1, 32–49 (2011).

16. Deb, K., Pratap, A., Agarwal, S. & Meyarivan, T. A fast and elitist multiobjective genetic algorithm: NSGA-II. IEEE transactions on evolutionary computation 6, 182–197 (2002).

17. Trinh, C. T., Liu, Y. & Conner, D. J. Rational design of efficient modular cells. Metabolic engineering 32, 220–231 (2015).

18. Burgard, A. P., Pharkya, P. & Maranas, C. D. Optknock: a bilevel programming framework for identifying gene knockout strategies for microbial strain optimization. Biotechnology and bioengineering 84, 647–657 (2003).

19. Klamt, S. & Mahadevan, R. On the feasibility of growth-coupled product synthesis in microbial strains. Metabolic engineering 30, 166–178 (2015).

20. Palsson, B. Ø. Systems biology: constraint-based reconstruction and analysis (Cambridge University Press, United Kingdom, 2015).

21. Feist, A. M. et al. Model-driven evaluation of the production potential for growth-coupled products of Escherichia coli. Metabolic engineering 12, 173–186 (2010).

22. Maranas, C. D. & Zomorrodi, A. R. Optimization Methods in Metabolic Networks (John Wiley & Sons, Hoboken, New Jersey, 2016).

23. Marler, R. T. & Arora, J. S. Survey of multi-objective optimization methods for engineering. Structural and multidisciplinary optimization 26, 369–395 (2004).

24. Monk, J. M. et al. iML1515, a knowledgebase that computes Escherichia coli traits. Nature biotechnology 35, 904 (2017).

25. Akita, H., Nakashima, N. & Hoshino, T. Pyruvate production using engineered Escherichia coli. AMB Express 6, 94 (2016).

26. Atsumi, S., Hanai, T. & Liao, J. C. Non-fermentative pathways for synthesis of branched-chain higher alcohols as biofuels. Nature 451, 86 (2008).

27. Layton, D. S. & Trinh, C. T. Engineering modular ester fermentative pathways in Escherichia coli. Metabolic Engineering 26, 77–88 (2014).

28. Niu, D. et al. Highly efficient L-lactate production using engineered Escherichia coli with dissimilar temperature optima for L-lactate formation and cell growth. Microbial cell factories 13, 78 (2014).

29. Rodriguez, G. M., Tashiro, Y. & Atsumi, S. Expanding ester biosynthesis in Escherichia coli. Nature Chemical Biology 10, 259–265 (2014).

30. Shen, C. R. et al. Driving Forces Enable High-Titer Anaerobic 1-Butanol Synthesis in Escherichia coli. Applied and Environmental Microbiology 77, 2905–2915 (2011).

31. Trinh, C. T., Unrean, P. & Srienc, F. Minimal Escherichia coli Cell for the Most Efficient Production of Ethanol from Hexoses and Pentoses. Applied and Environmental Microbiology 74, 3634–3643 (2008).

32. Tseng, H.-C. & Prather, K. L. Controlled biosynthesis of odd-chain fuels and chemicals via engineered modular metabolic pathways. Proceedings of the National Academy of Sciences, 201209002 (2012).

33. Yim, H. et al. Metabolic engineering of Escherichia coli for direct production of 1, 4-butanediol. Nature chemical biology 7, 445–452 (2011).

34. Yu, J., Xia, X., Zhong, J. & Qian, Z. Direct biosynthesis of adipic acid from a synthetic pathway in recombinant Escherichia coli. Biotechnology and bioengineering 111, 2580–2586 (2014).

35. Von Kamp, A. & Klamt, S. Growth-coupled overproduction is feasible for almost all metabolites in five major production organisms. Nature communications 8, 15956 (2017).

36. Hart, W. E. et al. Pyomo — Optimization Modeling in Python (Springer International Publishing, Cham, 2017).

37. Machado, D. & Herrgård, M. Systematic evaluation of methods for integration of transcriptomic data into constraint-based models of metabolism. PLoS Comput Biol 10, e1003580 (2014).

38. Lewis, N. E. et al. Omic data from evolved E. coli are consistent with computed optimal growth from genome-scale models. Molecular systems biology 6, 390 (2010).

39. Nocedal, J. & Wright, S. Numerical optimization (Springer Science & Business Media, United States of America, 2006).

40. Kaufman, D. E. & Smith, R. L. Direction choice for accelerated convergence in hit-and-run sampling. Operations Research 46, 84–95 (1998).

41. Heirendt, L. et al. Creation and analysis of biochemical constraint-based models: the COBRA Toolbox v3. 0. arXiv preprint arXiv:1710.04038 (2017).

42. King, Z. A. et al. Escher: a web application for building, sharing, and embedding data-rich visualizations of biological pathways. PLoS computational biology 11, e1004321 (2015).

43. Geoffrion, A. M. Generalized benders decomposition. Journal of optimization theory and applications 10, 237–260 (1972).

44. Fischetti, M., Ljubic, I. & Sinnl, M. Benders decomposition without separability: A computational study for capacitated facility location problems. European Journal of Operational Research 253, 557–569 (2016).

45. Von Kamp, A. & Klamt, S. Enumeration of Smallest Intervention Strategies in GenomeScale Metabolic Networks. PLOS Computational Biology 10, 1–13 (2014).

46. King, Z. A., O’Brien, E. J., Feist, A. M. & Palsson, B. O. Literature mining supports a next-generation modeling approach to predict cellular byproduct secretion. Metabolic Engineering 39, 220–227 (2017).

47. Fong, S. S. et al. In silico design and adaptive evolution of Escherichia coli for production of lactic acid. Biotechnology and bioengineering 91, 643–648 (2005).

48. Trinh, C. & Srienc, F. Metabolic engineering of Escherichia coli for efficient conversion of glycerol to ethanol. Appl Environ Microbiol 75, 6696–6705 (2009).

49. Garst, A. D. et al. Genome-wide mapping of mutations at single-nucleotide resolution for protein, metabolic and genome engineering. Nature biotechnology 35, 48 (2017).

50. Nielsen, D. R. et al. Engineering alternative butanol production platforms in heterologous bacteria. Metabolic engineering 11, 262–273 (2009).

51. Shi, A., Zhu, X., Lu, J., Zhang, X. & Ma, Y. Activating transhydrogenase and NAD kinase in combination for improving isobutanol production. Metabolic engineering 16, 1–10 (2013).

52. Hädicke, O., Bettenbrock, K. & Klamt, S. Enforced ATP futile cycling increases specific productivity and yield of anaerobic lactate production in Escherichia coli. Biotechnology and bioengineering 112, 2195–2199 (2015).

53. Khodayari, A. & Maranas, C. D. A genome-scale Escherichia coli kinetic metabolic model k-ecoli457 satisfying flux data for multiple mutant strains. Nature Communications 7 (2016).

54. Ishii, N. et al. Multiple high-throughput analyses monitor the response of E. coli to perturbations. Science 316, 593–597 (2007).

55. Kabir, M. M., Ho, P. Y. & Shimizu, K. Effect of ldhA gene deletion on the metabolism of Escherichia coli based on gene expression, enzyme activities, intracellular metabolite concentrations, and metabolic flux distribution. Biochemical Engineering Journal 26, 1–11 (2005).

56. Zhao, J., Baba, T., Mori, H. & Shimizu, K. Global metabolic response of Escherichia coli to gnd or zwf gene-knockout, based on 13 C-labeling experiments and the measurement of enzyme activities. Applied microbiology and biotechnology 64, 91–98 (2004).

57. Zhao, J. & Shimizu, K. Metabolic flux analysis of Escherichia coli K12 grown on 13C-labeled acetate and glucose using GC-MS and powerful flux calculation method. Journal of biotechnology 101, 101–117 (2003).

58. Zhou, S., Causey, T., Hasona, A., Shanmugam, K. & Ingram, L. Production of optically pure D-lactic acid in mineral salts medium by metabolically engineered Escherichia coli W3110. Appl. Environ. Microbiol. 69, 399–407 (2003).

59. Causey, T., Shanmugam, K., Yomano, L. & Ingram, L. Engineering Escherichia coli for efficient conversion of glucose to pyruvate. Proceedings of the National Academy of Sciences 101, 2235–2240 (2004).

60. Peng, L., Arauzo-Bravo, M. J. & Shimizu, K. Metabolic flux analysis for a ppc mutant Escherichia coli based on 13C-labelling experiments together with enzyme activity assays and intracellular metabolite measurements. FEMS Microbiology Letters 235, 17–23 (2004).

61. Siddiquee, K. A. Z., Arauzo-Bravo, M. & Shimizu, K. Metabolic flux analysis of pykF gene knockout Escherichia coli based on 13 C-labeling experiments together with measurements of enzyme activities and intracellular metabolite concentrations. Applied microbiology and biotechnology 63, 407–417 (2004).

62. Kim, P., Laivenieks, M., Vieille, C. & Zeikus, J. G. Effect of overexpression of Actinobacillus succinogenes phosphoenolpyruvate carboxykinase on succinate production in Escherichia coli. Appl. Environ. Microbiol. 70, 1238–1241 (2004).

63. Lin, H., Vadali, R. V., Bennett, G. N. & San, K.-Y. Increasing the acetyl-CoA pool in the presence of overexpressed phosphoenolpyruvate carboxylase or pyruvate carboxylase enhances succinate production in Escherichia coli. Biotechnology progress 20, 1599–1604 (2004).

64. Ravasz, E., Somera, A. L., Mongru, D. A., Oltvai, Z. N. & Barabási, A.-L. Hierarchical organization of modularity in metabolic networks. Science 297, 1551–1555 (2002).

65. Noor, E., Eden, E., Milo, R. & Alon, U. Central carbon metabolism as a minimal biochemical walk between precursors for biomass and energy. Molecular cell 39, 809–820 (2010).

66. Machado, D. & Herrgård, M. J. Co-evolution of strain design methods based on flux balance and elementary mode analysis. Metabolic Engineering Communications 2, 85–92 (2015).

67. Pharkya, P., Burgard, A. P. & Maranas, C. D. OptStrain: a computational framework for redesign of microbial production systems. Genome research 14, 2367–2376 (2004).

68. Pharkya, P. & Maranas, C. D. An optimization framework for identifying reaction activation/inhibition or elimination candidates for overproduction in microbial systems. Metabolic engineering 8, 1–13 (2006).

69. Chowdhury, A., Zomorrodi, A. R. & Maranas, C. D. k-OptForce: integrating kinetics with flux balance analysis for strain design. PLoS computational biology 10, e1003487 (2014).

70. Dinh, H. V., King, Z. A., Palsson, B. O. & Feist, A. M. Identification of growth-coupled production strains considering protein costs and kinetic variability. Metabolic engineering communications 7, e00080 (2018).

71. De Lorenzo, V. Systems biology approaches to bioremediation. Current opinion in biotechnology 19, 579–589 (2008).

72. Meyer, A. J., Segall-Shapiro, T. H., Glassey, E., Zhang, J. & Voigt, C. A. Escherichia coli “Marionette” strains with 12 highly optimized small-molecule sensors. Nature chemical biology, 1 (2018).

73. Fernandez-Rodriguez, J., Moser, F., Song, M. & Voigt, C. A. Engineering RGB color vision into Escherichia coli. Nature chemical biology 13, 706 (2017).

74. King, Z. A. et al. BiGG Models: A platform for integrating, standardizing and sharing genome-scale models. Nucleic acids research 44, D515–D522 (2015).

