## Supplementary Material SM2 for "Harnessing natural modularity of cellular metabolism to design a modular chassis cell for a diverse class of products by using goal attainment optimization": README.md

This file contains the release of ModCell2 used to generate the results of the following study:
- Title: Harnessing natural modularity of cellular metabolism to design a modular chassis cell for a diverse class of products by using goal attainment optimization
- Authors: Sergio Garcia and Cong Trinh
- Contact:

We recommend visiting the [online repository](https://github.com/TrinhLab/ModCell2) for the most updated versions of the software. Note that the sampling data for the genome-scale model is only [provided online](https://github.com/TrinhLab/modcell2-milp-sample-data) due to its large size.
